# Dual inhibition of lactate transporters MCT1 and MCT4 in pancreatic neuroendocrine tumors targets metabolic heterogeneity and functional redundancy

**DOI:** 10.1101/2025.01.31.635625

**Authors:** Konstantin Bräutigam, Janine Straub, Abdulloh K. Bihi, Renaud S. Maire, Simona Avanthay, Kristina Fillipova, Valentina Andreasi, Anna Battistella, Philipp P. Kirchner, Matthias S. Dettmer, Jörg Schrader, Marco Schiavo Lena, Stefano Partelli, Massimo Falconi, Ilaria Marinoni, Martin C. Sadowski, Aurel Perren

**Affiliations:** Institute of Tissue Medicine and Pathology, University of Bern, Bern, Switzerland; School of Medicine, Vita-Salute San Raffaele University, Milan, Italy; Pancreas Translational and Clinical Research Centre, Pancreatic Surgery Unit, IRCCS San Raffaele Scientific Institute, Milan, Italy; Department of Medicine, University Medical Center Hamburg-Eppendorf, Hamburg, Germany; Department of Pathology, IRCCS San Raffaele Scientific Institute, Milan, Italy

**Keywords:** pancreas, neuroendocrine, metabolic heterogeneity, monocarboxylate transporter (MCT), syrosingopine, tumoroids, adaptive metabolic reprogramming

## Abstract

Our current understanding of the metabolic landscape of pancreatic neuroendocrine tumors (PanNETs) is very limited. Such knowledge could lead the development of novel therapeutic strategies for subgroups of PanNET patients based on the metabolic profile of their tumor. Here, we investigated the expression of lactate transporters MCT1 and MCT4 in two independent PanNET cohorts (n=93; n=70) and analyzed their association with tumor aggressiveness and therapeutic vulnerability in cell lines, spheroids and patient-derived tumoroids of PanNET. Immunohistochemistry revealed four expressor types: MCT1/4-negative, MCT1-positive, MCT4-positive, and MCT1/4-double positive with frequent regional co-expression. Both homogenous and heterogenous expression patterns were observed, indicating metabolic heterogeneity within the latter subset of PanNETs. MCT4 expression correlated with the hypoxia marker CA9, suggesting a hypoxic and acidic tumor microenvironment.

Mechanistic studies revealed that MCT1 and MCT4 operate both as lactate efflux systems in PanNET cell lines, providing functional redundancy to their glycolytic roles. Inhibition of lactate efflux in normoxia and hypoxia using the dual MCT1/4 inhibitor syrosingopine significantly impaired lactate secretion, glycolysis, and proliferation across PanNET cell lines and 3D spheroid and patient-derived tumoroid models. In contrast, selective MCT1 or MCT4 inhibitors showed limited efficacy, underscoring the therapeutic need for co-targeting MCT1 and MCT4 due to functional redundancy and heterogenous expression. This work demonstrates MCT1 and MCT4 as metabolic markers and promising therapeutic targets of a subset of PanNETs with clinical features of aggressiveness.

## Introduction

Pancreatic Neuroendocrine Tumors (PanNETs) originate from the islets of Langerhans and represent a rare subtype (1-2%) of pancreatic neoplasms. PanNETs are broadly classified as ^··^functional^··^ when they secrete hormones or ^··^non-functional^··^ when having lost this ability. ∼80% of PanNETs belong to the non-functional category. The malignant potential of PanNETs increases with tumor size and grade. The mechanism of PanNET progression is complex and only partially understood due the clinical and genetic heterogeneity of this neoplasm. Frequent acquisition of mutations in epigenetic regulators MEN1, DAXX or ATRX (Marinoni et al. 2014; Singhi et al. 2017; Chou et al. 2018) in ∼40% of PanNET patients indicate that epigenetic changes play a critical role during tumor progression (Di Domenico et al. 2017). Molecular profiling approaches by genomics, transcriptomics and epigenomics have defined clinically and biologically distinct PanNET subtypes (Scarpa et al. 2017; Sadanandam et al. 2015; Di Domenico et al. 2020). Metabolic changes such as hypoxia and HIF-1α signaling were part of the molecular profiles of more aggressive PanNET subtypes (Scarpa et al. 2017; Sadanandam et al. 2015). Histological and molecular markers of hypoxia, such as low microvessel density and HIF-1α-regulated expression of carbonic anhydrase 9 (CA9) and glucose transporter 1 (GLUT1/SLC2A1) confirmed a link between hypoxia and clinico-pathological parameters of aggressive PanNET (Battistella et al., 2022; Couvelard et al., 2005). Furthermore, recent integrated transcriptomic and epigenomic analyses identified, among CA9 and GLUT1, four genes involved in lactate transport [monocarboxylate transporters MCT1 (SLC16A1) and MCT4 (SLC16A3)] and lactate metabolism (lactate dehydrogenases (LDH) LDHA and LDHB) as top upregulated genes in an aggressive subtype of primary PanNETs projected at the late stage of progression and termed ADM3 (Marinoni pers. communications). Like CA9 and GLUT1, MCT4 and LDHA are known transcriptional targets of HIF-1α and critical mediators of hypoxia-induced glycolysis and lactate metabolism (Ullah, Davies, and Halestrap 2006). As increased lactate transport is a characteristic of a high number of metastatic PanNET and seem epigenetically imprinted, dysregulated lactate transport may be a therapeutic target. Up to now, the functional role of dysregulated lactate metabolism and its potential as a therapeutic target have not been studied in human PanNET. Disrupting lactate transport via MCT1 or MCT4 inhibition has shown promise in preclinical studies (Baek et al., 2014) and early-phase clinical trials (Halford et al., 2023). However, metabolic tumor heterogeneity, metabolic symbiosis between tumor cells (Allen et al., 2016) or tumor cells and the stroma (Curry et al., 2013), and frequent co-expression of both transporters in tumors (Wang et al. 2021) suggest that combined inhibition of both lactate shuttles may be more effective in disrupting tumor growth (Sheng et al. 2023). Furthermore, compensatory MCT4 activity has been described as a resistance mechanisms to MCT1 inhibition (Le Floch et al., 2011; Polański et al., 2014).

MCT1 and MCT4 are proton-linked bidirectional monocarboxylate transporters (MCTs) localized at the plasma membrane. The direction of transport (uptake or efflux) of MCT1 and MCT4 is directed by their substrate affinity and its concentration gradient (Payen et al., 2020; Sharma et al., 2022). Despite their ability to transport similar cargos (pyruvate, lactate, ketone bodies such as acetoacetate and β-hydroxybutyrate, and short-chain fatty acids such as acetate, propionate, and butyrate), MCT1’s and MCT4’s affinities for these substrates are very different (Sharma et al., 2022). MCT4 almost exclusively operates as a lactate exporter, whereas MCT1 can operate bidirectionally to support a glycolytic phenotype by exporting lactate and pyruvate (Doherty et al., 2014), and an oxidative phenotype by importing lactate and pyruvate for mitochondrial respiration. This functional heterogeneity of MCT1 is often dictated by the tumor microenvironment (levels of oxygen, glucose and monocarboxylates) and can occur within different regions of the same tumor.

MCT1 and MCT4 are frequently overexpressed in various types of cancer, and their upregulation is correlated with a poor prognosis (Choi et al., 2014; Kim et al., 2015; Payen et al., 2020).

Several selective small molecule inhibitors against MCT1 and MCT4 have been developed (Wang et al. 2021), with AZD3965 showing promising anti-tumor activity in preclinical and clinical trials (Halford et al., 2023; Koltai and Fliegel, 2024). AZD3965 is a dual inhibitor of MCT1 (Ki=2 nM) and MCT2 (Ki=20 nM) without activity against MCT4 (Payen et al., 2020). AZD0095 is a recently developed selective inhibitor for MCT4 (Ki=1.3 nM) (Goldberg et al., 2023), and the repurposed anti-hypertensive drug syrosingopine is a dual MCT1 (IC_50_=2.5 μM) and MCT4 (IC_50_=40 nM) inhibitor but with a 60-fold higher activity against the latter transporter (Benjamin et al. 2018). Notably, MCT1 and MCT4 have not been investigated or pharmacologically targeted in PanNET, and their therapeutic potential remains to be elucidated.

While our current understanding of metabolic reprogramming in PanNET remains very limited, previous work strongly suggests the presence of metabolic subtypes in human PanNET (Sadanandam et al., 2015) involving hypoxia signalling. Here, we report on the classification of primary PanNET into four metabolic subtypes based on the expression of MCT1 and MCT4. Mechanistic studies of targeting MCT1 and MCT4 with proven inhibitors in 2D and 3D PanNET cell culture models, including patient-derived tumoroids evaluate their therapeutic potential.

## Materials and Methods

### Compounds

Syrosingopine (Sigma-Aldrich, SML1908-5MG), AZD0095 (MedChemExpress, HY-148517), AZD3965 (MedChemExpress, HY-12750), Sunitinib (Selleckchem, S1042), everolimus (Selleckchem, S1120) and SCH772984 (Selleckchem, S7101) were reconstituted in dimethylsulfoxide (DMSO) (Sigma-Aldrich, 472301). Metformin (Selleckchem, S1950) was dissolved in PBS. Stock solutions of compounds were stored at −20°C.

### Patient Cohorts

The study included 163 human tissue samples from two independent PanNET cohorts (n=93, University of Bern, Bern, Switzerland; n=70, San Raffaele, Milano, Italy; Table 1) with varying tumor grade, stage and intrapancreatic location. 22 (25.9%, Bern) and nine (13.0%, Milano) tumors were metastatic (M1) at diagnosis and initial resection of the primary tumor. Ki67 indices ranged from 0 to 40.0% (Milan: 0.6 to 65%) with a mean of 4.3% (Milano: 6.9%). Follow-up intervals, tumor recurrence and patient death were systematically recorded.

**Table 1.** Patient characteristics. *FU*: Follow-up. *DFS*: Disease Free Survival.

| Characteristic | Bern N = 93<sup>1</sup> |
| --- | --- |
| Age (at surgery) | 58.29 (14.16) |
| Female | 50 / 93 (54%) |
| Tumor Grade |  |
| 1 | 46 / 93 (49%) |
| 2 | 45 / 93 (48%) |
| 3 | 2 / 93 (2.2%) |
| Tumor localization |  |
| body | 17 / 93 (18%) |
| head | 34 / 93 (37%) |
| (Not specified) | 8 / 93 (8.6%) |
| tail | 34 / 93 (37%) |
| Tumor size (mm) | 34.32 (30.22) |
| Ki67 (%) | 4.28 (5.94) |
| (Missing) | 15 |
| recurrence | 19 / 78 (24%) |
| (Missing) | 15 |
| Time to recurrence (or follow-up) | 60.60 (56.78) |
| (Missing) | 13 |
| T-stage |  |
| 1 | 42 / 90 (47%) |
| 2 | 19 / 90 (21%) |
| 3 | 28 / 90 (31%) |
| 4 | 1 / 90 (1.1%) |
| (Missing) | 3 |
| N1 | 26 / 72 (36%) |
| (Missing) | 21 |
| M1 | 22 / 85 (26%) |
| (Missing) | 8 |
| DAXX |  |
| loss | 11 / 84 (13%) |
| retained | 72 / 84 (86%) |
| (Missing) | 10 |
| ATRX retained | 73 / 84 (87%) |
| (Missing) | 9 |

<sup>1</sup>n / N (%); Mean (SD)

| Characteristic | Milano N = 70<sup>1</sup> |
| --- | --- |
| Age (at surgery) | 56.37 (13.32) |
| Female | 34 / 70 (49%) |
| Tumor Grade |  |
| 1 | 25 / 70 (36%) |
| 2 | 41 / 70 (59%) |
| 3 | 4 / 70 (5.7%) |
| Tumor localization |  |
| body | 21 / 70 (30%) |
| body-tail | 2 / 70 (2.9%) |
| head | 28 / 70 (40%) |
| head-neck | 1 / 70 (1.4%) |
| neck | 1 / 70 (1.4%) |
| tail | 15 / 70 (21%) |
| uncinate process | 2 / 70 (2.9%) |
| Tumor size (mm) | 35.94 (20.92) |
| (Missing) | 1 |
| Ki67(%) | 6.94 (9.77) |
| T |  |
| 1 | 11 / 70 (16%) |
| 2 | 32 / 70 (46%) |
| 3 | 26 / 70 (37%) |
| 4 | 1 / 70 (1.4%) |
| N1 | 34 / 69 (49%) |
| (Missing) | 1 |
| M1 | 9 / 70 (13%) |
| DAXX loss | 19 / 70 (27%) |
| ATRX loss | 7 / 66 (11%) |
| (Missing) | 4 |
| recurrence | 21 / 70 (30%) |
| FU (months) | 55.49 (19.74) |
| DFS (months) | 44.64 (24.95) |
| Death | 5 / 70 (7.1%) |

<sup>1</sup>Mean (SD); n / N (%)

This study was approved by cantonal authorities (Kantonale Ethikkommission Bern, Ref.-Nr. KEK-BE 105/2015) in accordance with the Swiss Federal Human Research Act and the Italian ethic commission (Comitato Etico, CE 252/2019).

### Immunohistochemistry (IHC) and tissue microarrays (TMAs)

For each specimen, two (tumor center (TC) and front (TF)) representative tissue cores of 0.6-mm diameter were assembled into TMAs using a Grand Master automated tissue microarrayer (3DHistech, Budapest, Hungary). Sections (2.5□μm) were deparaffinized in a dewax solution (Leica Biosystems, Wetzlar, Germany) and rehydrated. Single-stain IHC was performed on a BOND RX automated immunostainer (Leica).

Primary antibodies were incubated for 30 min (MCT1, CA9, CD34) and 120 min (MCT4) at room temperature and used as follows: MCT1 (AB3538P, rabbit polyclonal; Merck, Darmstadt, Germany); dilution: 1:50; retrieval: EDTA, 40 min, 100°C; MCT4 (sc-376140, mouse monoclonal; Santa Cruz, Texas, USA); dilution: 1:50; retrieval: EDTA, 30 min, 95°C); Carboanhydrase 9 (CA9) (ab15086, rabbit polyclonal; Abcam, Cambridge, United Kingdom); dilution: 1:1500; retrieval: Tris EDTA, 30 min, 95°C, CD34 (#134M-16, mouse monoclonal; Cell Marque, CA, USA); dilution: 1:100; retrieval Tris EDTA, 30 min, 95°C.

Antibody detection was performed with the BOND Polymer Refine Detection kit (DS9800; Leica) using 3,3-diaminobenzidine (DAB) as a brown chromogen. The samples were counterstained with haematoxylin, dehydrated and mounted with Pertex (Sakura, Alphen aan den Rijn, the Netherlands). Slides were scanned on a Pannoramic 250 Flash scanner (3DHistech).

Staining was graded into three categories (negative, heterogeneously positive, and homogeneously positive) by four independent raters (KB, JS, VA, AP). Only membranous staining was considered positive, with heterogeneous (“het+”) expression defined as expression in 20-80% of tumor cells with a propensity to be near microvessels. Homogeneous (“hom+”) positivity was defined as homogenous expression in all tumor cells throughout the tissue core. In case of disagreement among raters, final consensus was reached on a multiheaded microscope. In case of divergent expression in TC and TF, the following rules were applied to assign a patient-level expression status: negative or homogeneous + heterogeneous = heterogeneous; homogeneous positive + negative = homogeneous. Microvessels were manually counted per tissue core in absolute numbers per mm² of tumor area using CD34 on our TMAs.

### Cell lines

The BON-1 cell line (RRID:CVCL_3985) was kindly provided by E.J.M. Speel. BON-1 cells were maintained in DMEM-F12 (10% FBS, 100□IU/mL penicillin, 0.1□mg/mL streptomycin). The QGP-1 cell line (RRID:CVCL_3143) was purchased from the Japanese Health Sciences Foundation. QGP-1 cells were cultured in RPMI 1640 medium (10% FBS, 100□IU/mL penicillin, 0.1□mg/mL streptomycin). NT-3, NT-18P and NT-18LM cells were kindly provided by J. Schrader and cultured in RPMI 1640 (10% FBS, 100□IU/mL penicillin, 0.1□mg/mL streptomycin) supplemented with human growth factors EGF (20□ng/mL, Gibco PHG0311) and bFGF (10□ng/mL, Gibco PHG0026) in collagen IV coated culture flasks. The mouse insulinoma PanNET cell line βTC-3 was kindly provided by R. Michael and cultured in high glucose (25 mM) DMEM (10% FBS, 2 mM glutamine, 100□IU/mL penicillin, 0.1□mg/mL streptomycin) For 3D cell culture, cells were seeded in 150□μL/well media (BON-1 300□cells/well, QGP-1 500□cells/well, NT-18P 1500□cells/well, NT-18LM 1500 cells/well, NT-3L1500 cells/well, and βTC-3 1000 cells/well) in a 96 well ultra-low attachment (ULA) round bottom BIOFLOAT plates (Sarstedt) and aggregated in the gravitational centre of the well by centrifugation (500□g for 10□min). Spheroid formation was continued by incubation for 3 days. For 2D and 3D cell culture, cells and spheroids were kept in a humidified incubator at 5% CO2 and 37□°C. For hypoxia experiments (1.5% O_2_), cells were transferred to a Whitley H35 HEPA hypoxystation (Don Whitley Scientific). 50% of volume was replenished with fresh media plus treatment after 4 days for long term treatments. Short tandem repeat (STR) analysis and regular mycoplasma testing by PCR confirmed the authenticity and health of the cell lines, respectively.

### PrestoBlue metabolic assay

For 2D experiments, BON-1 and QGP-1 (both 10,000 cells/well), βTC-3 (15,000 cells/well), NT-18P, NT-18LM and NT-3 (20,000 cell/well) were seeded in a 96 well plates in 150 μL of their respective growth media and cultured for 24 h before treatment. For better attachment of NT-18P, NT-18LM and NT-3 cells, wells were coated with collagen IV prior seeding. For 3D experiments, spheroids were generated as described above. Treatment of cells/spheroids was initiated by addition of 50 μL of growth media containing the indicated compounds at 4x final concentration. For hypoxia experiments, the treated samples were transferred to a Whitley H35 HEPA hypoxystation (Don Whitley Scientific), incubated for the indicated times at 1.5% O2, 5% CO2, and 37°C, and sample processing was carried out in the hypoxystation. At the indicated endpoint of treatment, the media volume was reduced to 90 μL/well, and 10 μL/well of the resazurin-based PrestoBlue reagent (Invitrogen, A13261) was added. After 45-60min incubation in a humidified incubator at 5% CO2 and 37□°C, the resazurin to resorufin conversion was measured in fluorescence mode with a 540 nm excitation and 610 nm emission setting on a microplate reader (TECAN Infinite 200 PRO). After background correction, the metabolic activity was calculated relative to the vehicle control (mean of n=3 ±STD).

### Lactate secretion assay

Cells were seeded as described for the PrestoBlue assay. Growth media was completely removed and replaced with serum-free media supplemented with the indicated treatments. For NT-18P, NT-18LM and NT-3 cells, serum-free media also contained EGF and bFGF. For hypoxia experiments, the treated samples were transferred to a Whitley H35 HEPA hypoxystation (Don Whitley Scientific), incubated for the indicated times at 1.5% O2, 5% CO2, and 37°C, and sample collection was carried out in the hypoxystation. At treatment endpoint, 100 μL of conditioned media was removed and cleared by centrifugation (5 min, 15,000 rcf). Lactate was quantified in dilutions of the supernatant (BON-1 1:150, QGP-1 1:100, NT-18P 1:50, NT-18LM 1:50 and NT-3 1:40) using the L-Lactate Assay Kit (Sigma-Aldrich, MAK443) with a modified protocol, i.e., the reaction volume was reduced from 100 μL to 30 μL with proportional adjustments of reagent and test sample volumes and processing in clear V-shaped bottom 96 well plates. After incubation in the dark at room temperature for one hour, fluorescence of the lactate reporter was measured using a microplate reader (TECAN Infinite 200 PRO) with an excitation/emission filter setting of 530 nm/585 nm. After background correction, linear regression analysis was performed to quantify lactate concentrations of the diluted samples and adjusted for the dilution factor (mean of n=3 ±STD).

### Seahorse XFe96 metabolic flux analysis

BON-1 (17,500 cells/well), QGP-1 (20,000 cells/well), NT-18P and NT-18LM cells (both 30,000 cells/well) were seeded in 100 μL/well of their respective growth media in Seahorse XFe96 microplates (Agilent) and incubated for 24 h. For better attachment of NT-18P and NT-18LM cells, wells were coated with collagen IV prior seeding. For analysis in hypoxia, XFe96 microplates were transferred 6 h post seeding to a Whitley H35 HEPA hypoxystation (Don Whitley Scientific), incubated for two days at 1.5% O2, 5% CO2, and 37°C, and sample processing and metabolic flux analysis were carried out in the hypoxystation. To run the glycolysis stress test, the medium was changed to basal serum-, glucose- and pyruvate-free medium (Seahorse XF RPMI 103576-100, Agilent) supplemented with 2 mM L-glutamine (Sigma Aldrich, #1294808) through repeated partial washing steps as per manufacturer’s instructions. The four injections ports were loaded with the following compounds to yield the treatment concentration as shown in brackets: A, indicated MCT1/2/4 inhibitors; B, glucose (5 mM, ThermoFisher, #A2494001); C, oligomycin (1.5 μM); D, 2-deoxyglucose (8.33 mM). After calibration of the Seahorse XFe96 Analyzer, a modified assay protocol was run to measure both oxygen consumption rate (OCR) and extracellular acidification rate (ECAR): 1) four baseline measurements, 2) injection from port A followed by four measurements, 3) injection from port B followed by four measurements, 4) injection from port C followed by four measurements, 5) injection from port D followed by five measurements. The interval between measurements was 5 min for all five analysis steps. Every data point is a representative of 5-6 technical replicates (mean ±STD). The ECAR and OCR measurements were analyzed using Wave desktop software (Agilent).

### RNA extraction and qRT-PCR

Approximately 5 x 10^5^ cells were briefly washed with ice cold PBS before RNA extraction commenced using the RNeasy® Mini Kit (Qiagen, 74104) according to the manufacturer’s instructions. Before elution, RNA was treated with DNase (Qiagen) to remove genomic DNA and improve RNA purity. Concentration of RNA was measured using a NanoDrop ND-1000 Spectrophotometer (ThermoScientific), and cDNA was prepared from 2 μg total RNA using the High Capacity cDNA Reverse Transcription Kit (Applied Biosystems, 4368814). qRT-PCR was performed with SYBR Green PCR Master Mix (Invitrogen) on a ViiA-7 Real-Time PCR system (Applied Biosystems). Determination of relative mRNA levels was calculated using the comparative ΔΔCT method compared to the expression of housekeeping gene Receptor-like protein 32 (RPL32) in each treatment and calculated as fold change relative to vehicle control (DMSO). All experiments were performed in triplicate, and analysis and statistics were performed with GraphPad Prism software. Primer sequences for MCT4 and RPL32 can be found in supplementary information (Table S7).

### Cell and spheroid growth analysis

Growth as a function of increasing cell confluence (2D) or spheroid size (3D) was measured by real-time microscopy in a humidified environment at 5% CO2 and 37□°C using the Cell-IQ v2 SLF system (CM Technologies). For 2D experiments, BON-1 (5000 cells/well), QGP-1 (6000 cells/well), NT-18P and NT-18LM cells (both 8000 cells/well) were seeded in 150 μL/well of their respective growth media in 96 microplates and incubated for 24 h. For better attachment of NT-18P and NT-18LM cells, wells were coated with collagen IV prior seeding. For 3D experiments, spheroids were generated in ULA plates as described above. After treatment, cell culture plates were transferred into the Cell-IQ systems and imaged at 10x magnification at the indicated time intervals. 2D cell proliferation was quantified by measuring cell confluence of one field-of-view per well over time using CellActivision Software (Yokogawa). For assessment of spheroid growth, a 3×3 grid (9 field of views) was imaged with the focus set to the capture the maximum diameter of the spheroids. Images were stitched using CellActivision Software (Yokogawa), and spheroid area was measured using CellProfiler Software 4.2.5 (Broad Institute) with a custom-made analysis pipeline. 2D and 3D growth curves were calculated by normalization of the measurements with the corresponding t=0 values of cell confluence and spheroid area, respectively.

### Primary PanNET cell isolation and tumoroid 3D culture

Isolated primary PanNET cells and tumoroids were cultured in AdvDMEM plus 5% FBS, Hepes 10□mM, 1% L-glutamine, 1% penicillin-streptomycin-amphotericin B and growth factors (20□ng/mL EGF, 10□ng/mL bFGF (Thermo Fisher Scientific, USA), 100□ng/mL PlGF, 769□ng/mL IGF-1 (Selleckchem, USA)) in 96 well ULA BIOFLOAT plates (Sarstedt). Patient-derived tumoroids (PDTs) were established from cryopreserved tumor tissues of primary (7) and metastatic (1) G1/G2 PanNETs as described previously(April-Monn et al. 2020; April-Monn et al. 2024). In brief, cryopreserved tissues were dissociated with a gentleMACS® dissociator (Myltenyi Biotec, Switzerland) in AdvDMEM medium, 0.25% Trypsin (Sigma-Aldrich, Switzerland), 10□mg/ml collagenase IV, (Worthington, USA), 10□mg/ml DNAse (Roche, Switzerland). After dissociation red blood cells were lysed for 3□min with ACK lysis buffer (Thermo Fisher Scientific, USA) at room temperature. Fibroblast cells were partially removed by attachment, involving a short incubating of the cell suspension at 37C for 1□h in coated plates. The fibroblast-depleted supernatant was collected and seeded on ULA plates to recover. After 2 days, the cell suspension was removed from the plates and centrifuged at 120□g for 5□min to separate cells and aggregates from debris/dead cells. The cell pellet was carefully resuspended in fresh AdvDMEM + growth factors medium and counted. For PDT formation, 3000-5000 cells/well were seeded in 50□μL/well AdvDMEM + growth factors medium supplemented with 123□μg/mL growth-factor-reduced Matrigel in 96-well ULA plates and centrifuged (500□g for 10□min). PDT formation was continued by incubation for 3 days before treatment with the indicated inhibitors. For validation of tumor cell content (synaptophysin-positive) and target expression (MCT1 and MCT4), micro cell blocks were prepared from untreated samples taken at day 0 of PDT seeding (single cell suspension) and at endpoint (tumoroid) and processed by immunohistochemistry as described previously (April-Monn et al. 2020).

### Real-time-Glo™ metabolic activity of PDTs

Real-Time-GloTM MT Cell Viability (RTG) assay (Promega, Switzerland) was used to continuously monitor metabolic activity (reductive power) of PDTs. The RTG assay was performed according to the manufacturer’s instructions, and luminescence was measured in an Infinite® 200 PRO plate reader (Tecan, Switzerland). On the day of treatment, conditioned medium of each well was supplemented with 50 μL of fresh AdvDMEM + GF medium containing Matrigel and RTG assay reagents to a final volume of 100 μL, and an RTG baseline luminescence reading was recorded 6 h prior drug treatment. After treatment start, RTG luminescence was recorded every 24 h or as indicated. Growth factors and FBS were replenished every 3–4 days. To calculate drug response curves, relative luminescence unit (RLU) readings of each treatment were calculated using their corresponding baseline values and then normalized to vehicle control (DMSO) using the same time point.

### Live cell fluorescent microscopy

For 2D experiments, BON-1 (10,000 cells/well), QGP-1 (10,000 cells/well), NT-18P and NT-18LM cells (both 20,000 cells/well) were seeded in 150 μL/well of their respective growth media in μ-Plate 96-well optical μ-Plates (IBIDI) and incubated for 24 h. For better attachment of NT-18P and NT-18LM cells, wells were coated with collagen IV prior seeding. For 3D live cell microscopy, cells were seeded in 150□μL/well media (BON-1 300□cells/well, QGP-1 500□cells/well, NT-18P 1500□cells/well, and NT-18LM 1500□cells/well) in a 96 well ultra-low attachment (ULA) round bottom BIOFLOAT plates (Sarstedt) as described above. After treatment for the indicated time, culture media was reduced to 90 μL/well, and 10 μL of a 10-fold concentrated staining mix was added to each well. For assessing live cell count (SytoxGreen-negative) and cell death (SytoxGreen-positive), cells were stained with 10 μM Hoechst 33342 (MedChemExpress) and 0.5 μM SytoxGreen (ThermoFisher Scientific). Mitochondrial mass (active and inactive) and activity were assessed by labelling with 150 nM MitoTracker Green (ThermoFisher Scientific) and 100 nM MitoTracker Orange CMTMRos (ThermoFisher Scientific). Lysosomes were stained with 30 nM LysoTracker Deep Red (ThermoFisher Scientific), and cellular ROS was visualized by 2.5 μM CellROX Deep Red (ThermoFisher Scientific). Uptake of glucose and fatty acids were assessed with 100 μM 2-NBDG (ThermoFisher Scientific) and 2.5 μM Bodipy FL C16 (ThermoFisher Scientific), respectively. Free cholesterol, neutral (lipid droplets) and phospholipid lipid content were labelled with 50 μg/mL filipin III (Sigma-Aldrich) and 100 ng/mL Nile Red (Sigma-Aldrich), respectively. For 2D experiments, cells were labelled by incubation in the dark for 60-120 min, whereas spheroids and tumoroids were labelled for 3-4 h. Fluorescence microscopy was carried out at the indicated magnification on IN Cell Analyzer 2000 (GE Healthcare) or CQ1 (Yokogawa) microscope system. For quantitative analysis of fluorometric and morphometric parameters, images were analyzed with custom-made pipelines using Cell Profiler software (Broad Institute).

### Statistical analysis

Statistical analyses were performed using GraphPad Prism version 10 (GraphPad Software, Inc, California, USA) and the R statistical computing environment version 4.4.1 (RStudio version 2024.04.2). All p-values were calculated for two-tailed tests with significance set at p<0.05. Statistical analyses included Student’s t-test and Fisher and Kruskal-Wallis tests for pairwise comparisons, one-way ANOVA for multiple comparisons (Bonferroni post-hoc analysis), and Kaplan-Meier survival analysis with log-rank tests for time-to-event data.

## Results

### Expression of MCT1 and MCT4 defines metabolic subgroups in PanNET

Immunohistochemistry of MCT1 and MCT4 with tissue microarrays of two independent PanNET cohorts (Bern and Milano, Table 1) showed positive plasma membrane expression in a subset of PanNET samples (Figure 1A). MCT1 expression was detected in 13% (12/92 Bern, Table 2) and 17% of patient samples (12/70 Milano, Table 2), and MCT4 expression in 17% (15/90 Bern, Table 2) and 30% of patient samples (21/70 Milano, Table 2). Notably, expression of the hypoxia marker CA9 correlated well with MCT4 expression in both cohorts (Spearman-ρ=0.65 Bern; Spearman-ρ=0.67 Milano). A subset of MCT1/4-positive PanNETs of both cohorts showed co-expression of both lactate shuttles (Figs. 1AB), often within the same tissue regions (SFig. 1). MCT1/4-positive PanNET tissues could be further classified based on the expression pattern of both transporters (Fig. 2A). We observed a slightly higher abundance of tissue cores with homogenous expression patterns (Table 2). Heterogeneous MCT4 PanNET cores often presented with congregated expression distant to microvessels, suggesting regional hypoxia. In contrast, homogenous MCT4 expression was present even in PanNET cores with high MVD (Fig. 2A), raising the possibility of pseudohypoxia by epigenetic imprinting in this subtype.

**Figure 1.**
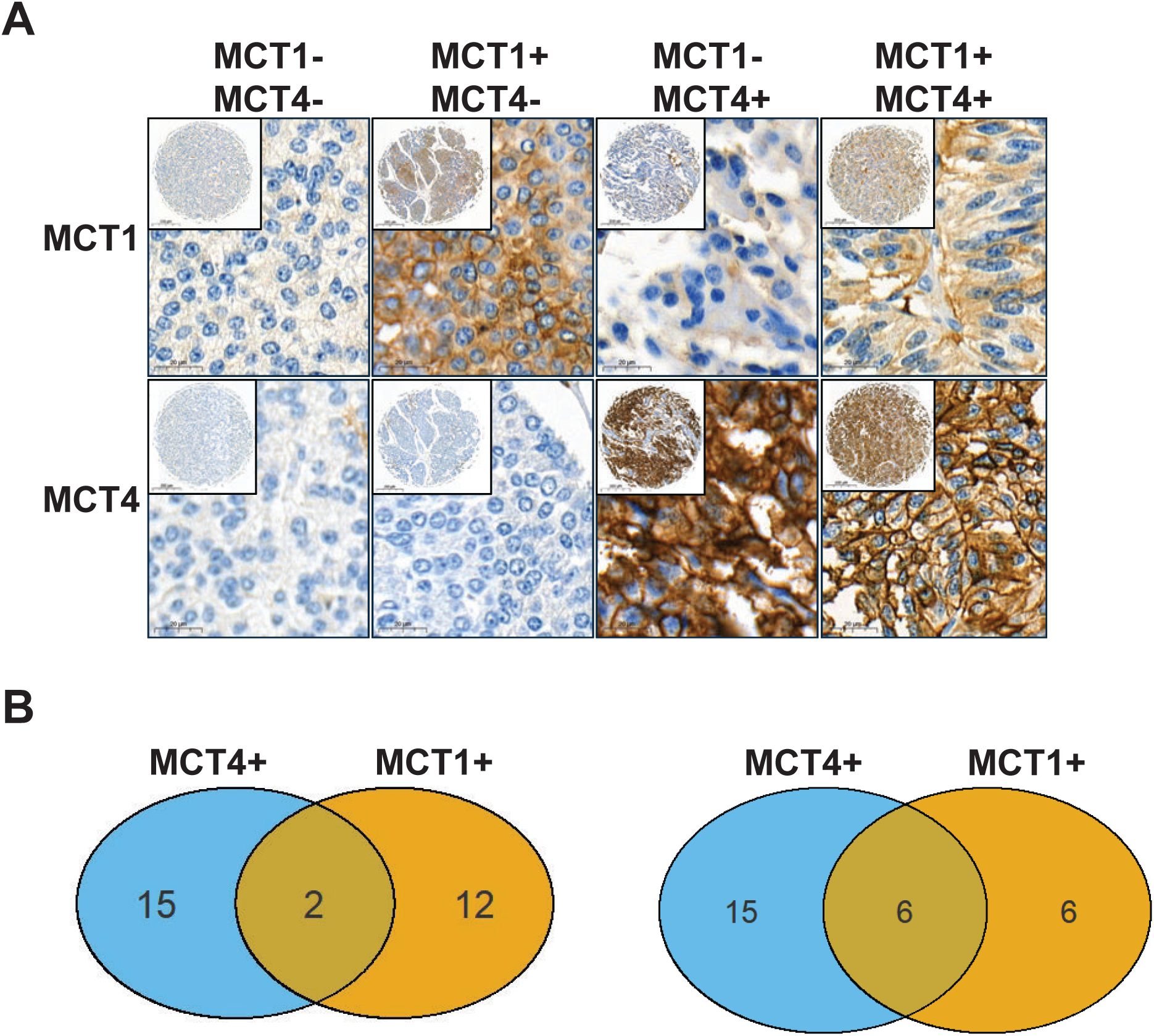
(A) Patterns of immunohistochemical membranous MCT1/MCT4 protein expression in PanNET. From left to right, double-negative (MCT1-/MCT4-), single-positive and double positive expression (MCT1+/MCT4+) in representative tumors. *Inset:* Overview of tissue core. (B) Venn diagrams of exclusive and co-expression of MCT1and MCT4 in the Bern (left panel) and Milano cohort (right panel). The majority of MCT-expressors of both cohorts was MCT4+/MCT1-.

**Figure 2.**
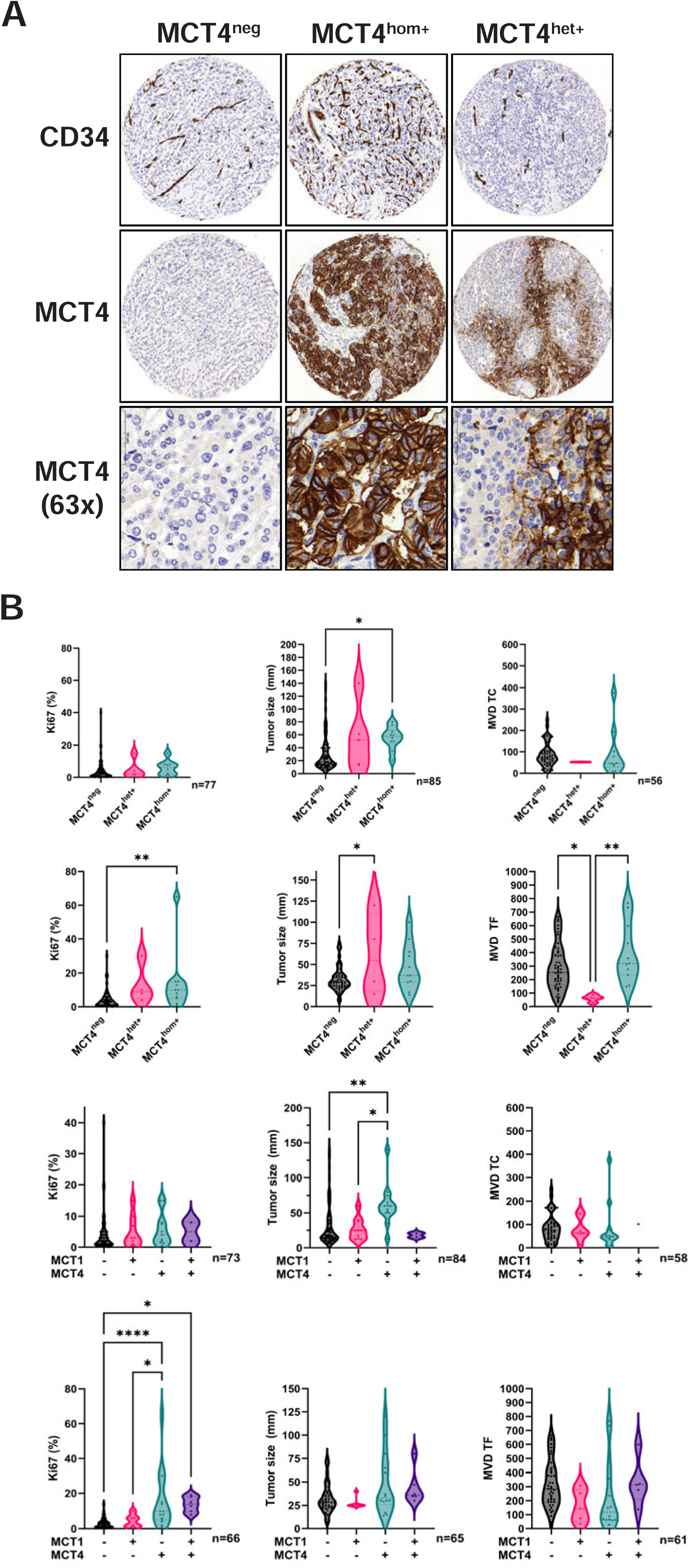
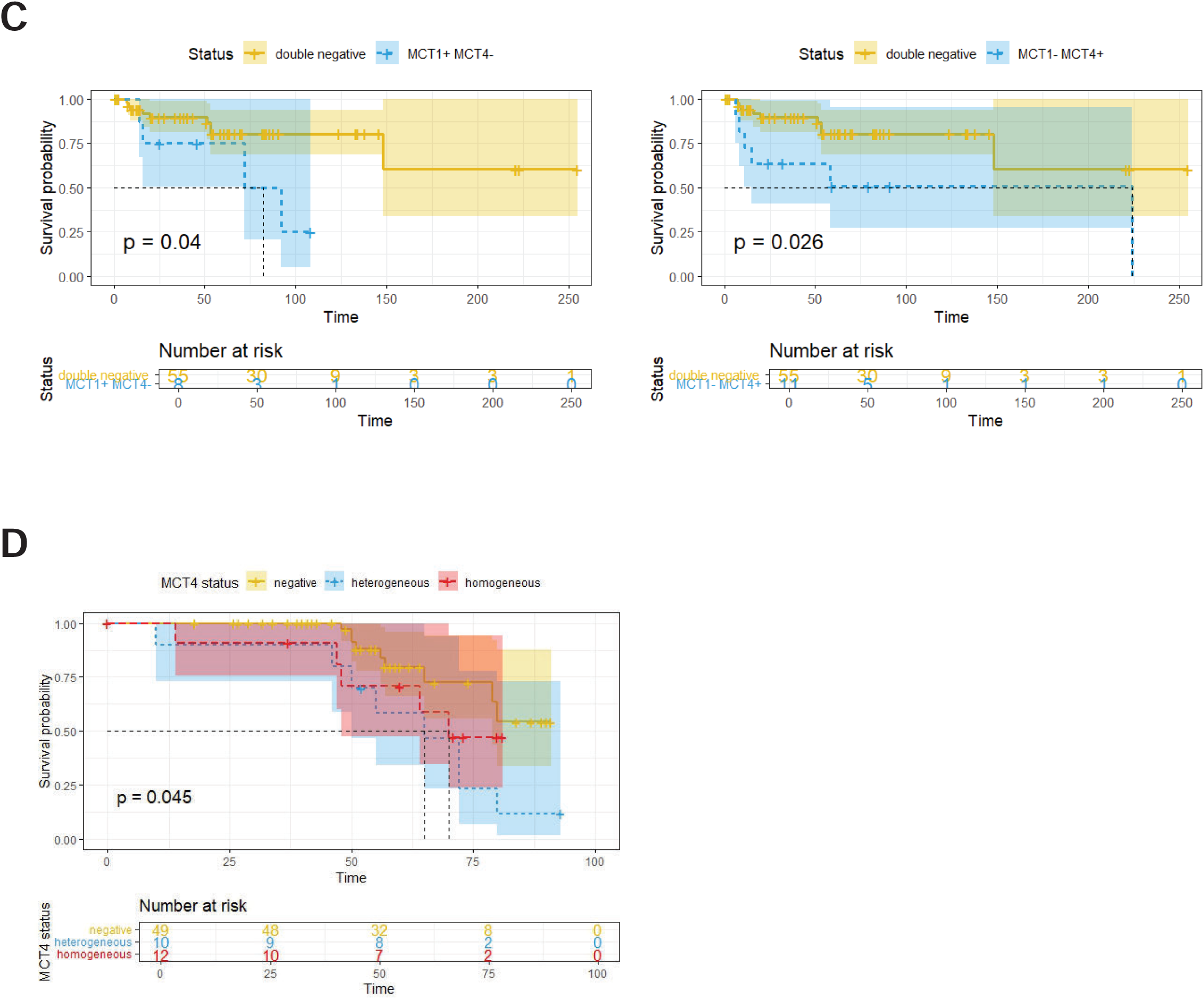
(A) Immunohistochemistry of CD34-positive microvessels and MCT4 expression in tissue cores of PanNET revealed distinct MCT4 expression patterns, indicating pseudohypoxia in MCT4 homogenous-positive tumors (MCT4^hom+^) and regional hypoxia in MCT4 heterogeneous-positive tumors (MCT4^het+^). (B) Violin plots of Ki67 proliferation index, tumor size, and microvessel density (MVD) of MCT4-negative, -heterogeneous-positive and homogeneous-positive PanNETs of the Bern (top panels) and Milano cohorts (bottom panels). MCT4 expressors associate with larger tumor size and higher Ki67 indices. MCT4 heterogeneous-positive tumors show less MVD. The same analysis is shown for double-negative, single and co-expressors for both cohorts. (C) Survival analysis (tumor relapse) differentiated by MCT1 and MCT4 status of the Bern cohort and (D) homogeneous and heterogenous expression patterns of the Bern (top panels) and Milano cohorts (bottom panel). Single-expressors (C) and MCT4-heterogeneous tumors (D) relapse significantly earlier than double-negative tumors. *TC*: Tumor Center; *TF*: Tumor Front.

**Table 2.**
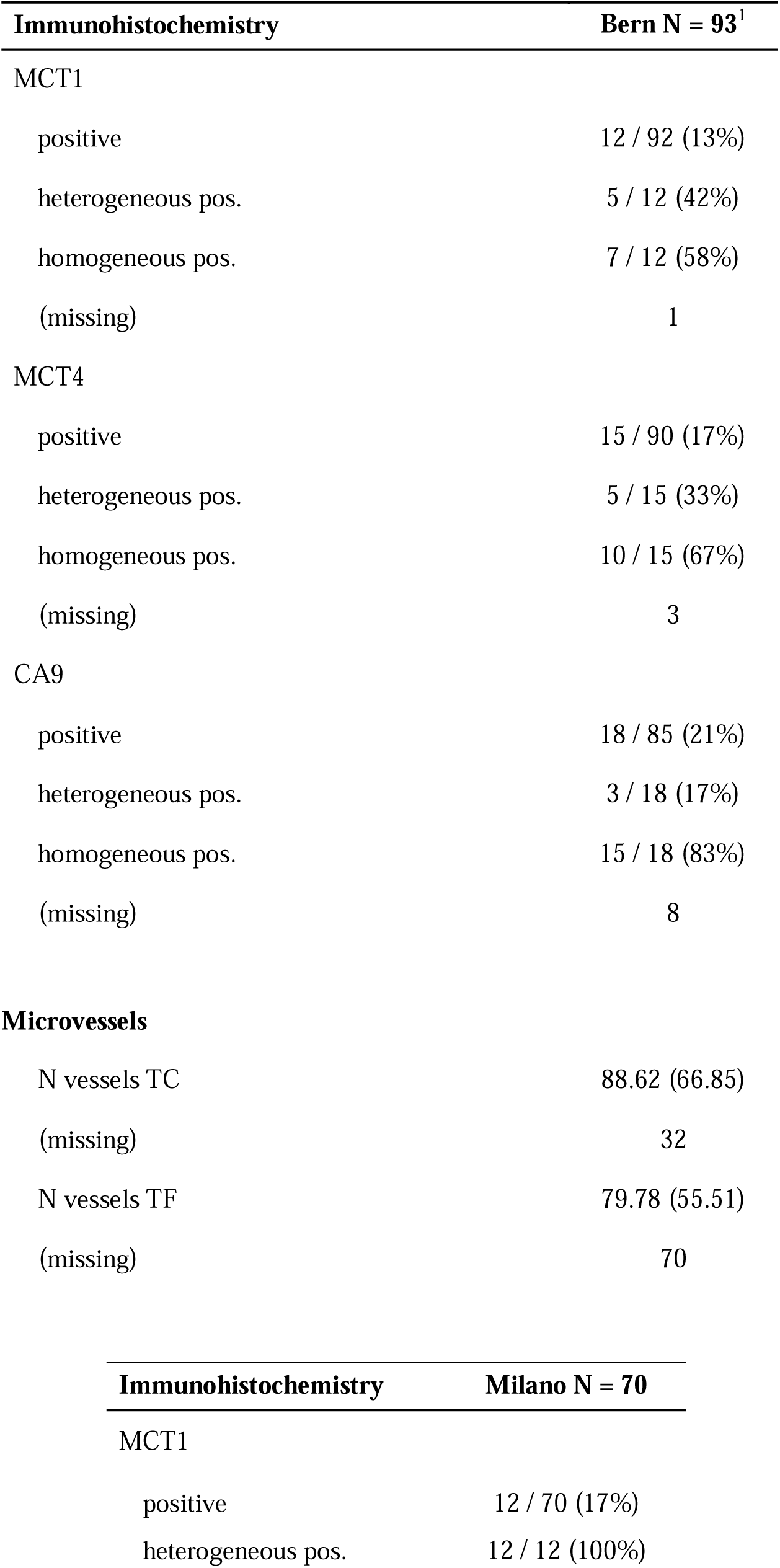
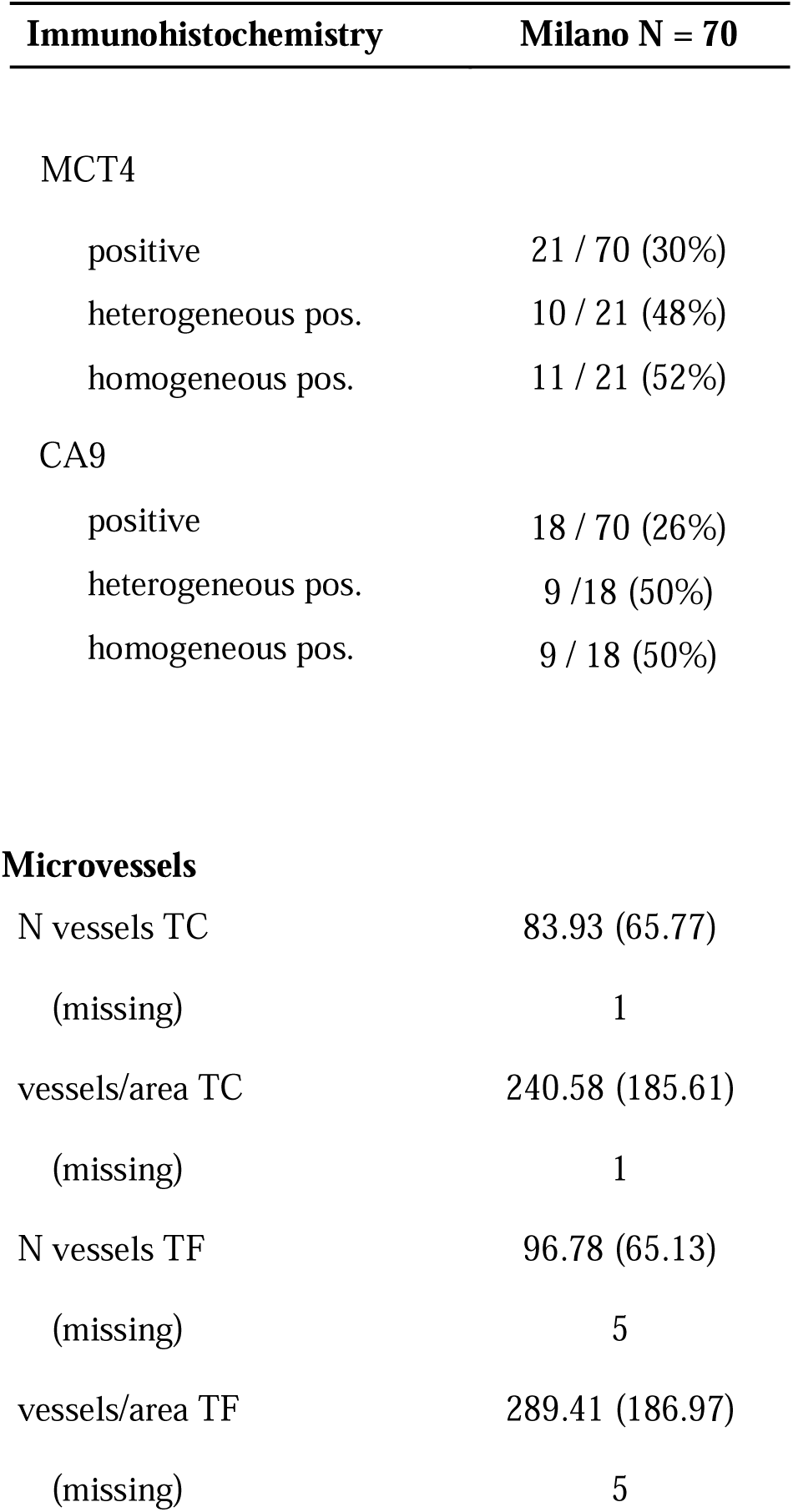
Immunohistochemistry and MVD. *TC*: Tumor Center. *TF*: Tumor Front.

| Immunohistochemistry | Bern N = 93<sup>1</sup> |
| --- | --- |
| MCT1 |  |
| positive | 12 / 92 (13%) |
| heterogeneous pos. | 5 / 12 (42%) |
| homogeneous pos. | 7 / 12 (58%) |
| (missing) | 1 |
| MCT4 |  |
| positive | 15 / 90 (17%) |
| heterogeneous pos. | 5 / 15 (33%) |
| homogeneous pos. | 10 / 15 (67%) |
| (missing) | 3 |
| CA9 |  |
| positive | 18 / 85 (21%) |
| heterogeneous pos. | 3 / 18 (17%) |
| homogeneous pos. | 15 / 18 (83%) |
| (missing) | 8 |
| **Microvessels** |  |
| N vessels TC | 88.62 (66.85) |
| (missing) | 32 |
| N vessels TF | 79.78 (55.51) |
| (missing) | 70 |

| Immunohistochemistry | Milano N = 70 |
| --- | --- |
| MCT1 |  |
| positive | 12 / 70 (17%) |
| heterogeneous pos. | 12 / 12 (100%) |

| Immunohistochemistry | Milano N = 70 |
| --- | --- |
| MCT4 |  |
| positive | 21 / 70 (30%) |
| heterogeneous pos. | 10 / 21 (48%) |
| homogeneous pos. | 11 / 21 (52%) |
| CA9 |  |
| positive | 18 / 70 (26%) |
| heterogeneous pos. | 9 / 18 (50%) |
| homogeneous pos. | 9 / 18 (50%) |
| Microvessels |  |
| N vessels TC | 83.93 (65.77) |
| (missing) | 1 |
| vessels/area TC | 240.58 (185.61) |
| (missing) | 1 |
| N vessels TF | 96.78 (65.13) |
| (missing) | 5 |
| vessels/area TF | 289.41 (186.97) |
| (missing) | 5 |

### MCT1 and MCT4 expression patterns correlate with adverse prognosis in PanNET

In the Bern cohort, MCT1 expression was significantly associated with M1-stage (p=0.024, Table 3). All (3/3) heterogeneous MCT1-positive PanNETs were M1 at resection. MCT4-positive tumors were significantly larger in size (p=0.003). MCT1/4-positive PanNETs (single and co-expressors) were more frequently of N1-stage (13/23, 56.5%) than MCT1/4-negative PanNETs (13/47, 27.7%). Exclusive MCT4 expressors (MCT1-negative) and homogeneous MCT4 expressors were significantly larger in size than MCT1/MCT4-double positive (p<0.05) and MCT4-negative PanNETs (p=0.003), respectively (Fig. 2B).

**Table 3.** Correlation analysis, Bern.

**MCT1**
| Clinical Variable | Statistical Test | P-value | Significant |
| --- | --- | --- | --- |
| gender | Fisher test | 0.124 | Not Significant |
| T | Fisher test | 0.514 | Not Significant |
| N | Fisher test | 0.226 | Not Significant |
| M | Fisher test | 0.024 | Significant |
| DAXX | Fisher test | 1.000 | Not Significant |
| ATRX | Fisher test | 1.000 | Not Significant |
| tumor size | Kruskal-Wallis | 0.625 | Not Significant |
| WHO Grading | Fisher test | 0.861 | Not Significant |

**MCT4**
| Clinical Variable | Statistical Test | P-value | Significant |
| --- | --- | --- | --- |
| gender | Fisher test | 0.124 | Not Significant |
| T | Fisher test | 0.514 | Not Significant |
| N | Fisher test | 0.226 | Not Significant |
| M | Fisher test | 0.024 | Significant |
| DAXX | Fisher test | 1.000 | Not Significant |
| ATRX | Fisher test | 1.000 | Not Significant |
| tumor size | Kruskal-Wallis | 0.003 | Significant |
| WHO Grading | Fisher test | 0.861 | Not Significant |

**MCT1/4 Single and Dual Expressors**
| Clinical Variable | Statistical Test | P-value | Significant |
| --- | --- | --- | --- |
| gender | Fisher test | 0.124 | Not Significant |
| T | Fisher test | 0.514 | Not Significant |
| N | Fisher test | 0.226 | Not Significant |
| M | Fisher test | 0.024 | Significant |
| DAXX | Fisher test | 1.000 | Not Significant |
| ATRX | Fisher test | 1.000 | Not Significant |
| tumor size | Kruskal-Wallis | 0.004 | Significant |
| WHO Grading | Fisher test | 0.861 | Not Significant |

In the Milano cohort, homogeneous MCT1 expression was significantly associated with higher Ki67 proliferation indices (p=0.038, Table 4 and SFig. 2A) and tended to have a positive nodal (N1) stage (p=0.06, Table 4). Expression of MCT4 significantly associated with higher T-stage (p=0.034) and higher Ki67 (p=0.002). Ki67 was significantly higher in single MCT4 and MCT1/MCT4 double expressors compared to non-expressors (Fig. 2B). Heterogeneous MCT4 tumors were larger in size and had a significantly lower microvessel density (MVD) than homogeneous or non-expressors (Fig. 2B).

**Table 4.** Correlation analysis, Milano.

**MCT1**
| Clinical Variable | Statistical Test | P-value | Significant |
| --- | --- | --- | --- |
| T | Fisher test | 0.239 | Not Significant |
| N | Fisher test | 0.062 | Not Significant |
| M | Fisher test | 0.177 | Not Significant |
| age | Kruskal-Wallis | 0.374 | Not Significant |
| gender | Fisher test | 0.213 | Not Significant |
| tumor size | Kruskal-Wallis | 0.606 | Not Significant |
| Ki67 | Kruskal-Wallis | 0.038 | Significant |
| DAXX | Fisher test | 0.491 | Not Significant |
| ATRX | Fisher test | 0.084 | Not Significant |
| recurrence | Fisher test | 0.489 | Not Significant |
| death | Fisher test | 1.000 | Not Significant |

**MCT4**
| Clinical Variable | Statistical Test | P-value | Significant |
| --- | --- | --- | --- |
| T | Fisher test | 0.034 | Significant |
| N | Fisher test | 1.000 | Not Significant |
| M | Fisher test | 0.169 | Not Significant |
| age | Kruskal-Wallis | 0.154 | Not Significant |
| gender | Fisher test | 0.221 | Not Significant |
| tumor size | Kruskal-Wallis | 0.164 | Not Significant |
| Ki67 | Kruskal-Wallis | 0.002 | Significant |
| DAXX | Fisher test | 0.116 | Not Significant |
| ATRX | Fisher test | 0.530 | Not Significant |
| recurrence | Fisher test | 0.068 | Not Significant |
| death | Fisher test | 0.126 | Not Significant |

**MCT1/4 Single and Dual Expressors**
| Clinical Variable | Statistical Test | P-value | Significant |
| --- | --- | --- | --- |
| T | Fisher test | 0.108 | Not Significant |
| N | Fisher test | 0.185 | Not Significant |
| M | Fisher test | 0.125 | Not Significant |
| age | Kruskal-Wallis | 0.360 | Not Significant |
| gender | Fisher test | 0.318 | Not Significant |
| tumor size | Kruskal-Wallis | 0.101 | Not Significant |
| Ki67 | Kruskal-Wallis | 0.000005 | Significant |
| DAXX | Fisher test | 0.307 | Not Significant |
| ATRX | Fisher test | 0.069 | Not Significant |
| recurrence | Fisher test | 0.011 | Significant |
| death | Fisher test | 0.072 | Not Significant |

Importantly, expression of either lactate shuttle was significantly correlated with earlier tumor relapse (MCT1 p=0.04, MCT4 p=0.026) when compared to negative PanNETs or MCT1/4 co-expressors in the Bern cohort (Table 3 and Fig 2C). PanNET patients with heterogenous MCT4 expression trended towards earlier relapse (p=0.23 Bern, p=0.045 Milano) in comparison to negative and homogenous-positive tumors (Fig. 2D).

Taken together, expression of MCT1 and MCT4 defines four metabolic subgroups (MCT1/4-double negative, MCT1+/MCT4-, MCT1-/MCT4+, MCT1/4-double positive) in both independent PanNET cohorts which correlated with clinico-pathological parameters of aggressive disease. The known bi-directional lactate transport capability of MCT1 and the strong association of MCT4 with hypoxia together with the observed expression patterns of both MCT1 and MCT4 suggest high metabolic heterogeneity within PanNET cores and across PanNET patients.

### PanNET cells secrete lactate and express MCT1 and MCT4

Lactate metabolism and the role of lactate transporters such as MCT1 and MCT4 have not been studied in human PanNET. As shown in Figure 3A, all human PanNET cell lines secreted lactate in hypoxia (1.5% O_2_, 48h), whereas only the NEC cell-lines BON-1 and QGP-1 secreted considerable amounts of lactate when cultured in normoxic conditions, indicating aerobic glycolysis (Warburg effect). Nuclear magnetic resonance analysis of glucose consumption and lactate secretion confirmed above observations (data not shown). The glycolytic phenotype of BON-1 and QGP-1 cells was consistent with a several-fold higher metabolic activity in normoxia and hypoxia when compared to NT-18P, NT-18LM and NT-3 cells (SFig. 3A).

**Figure 3.**
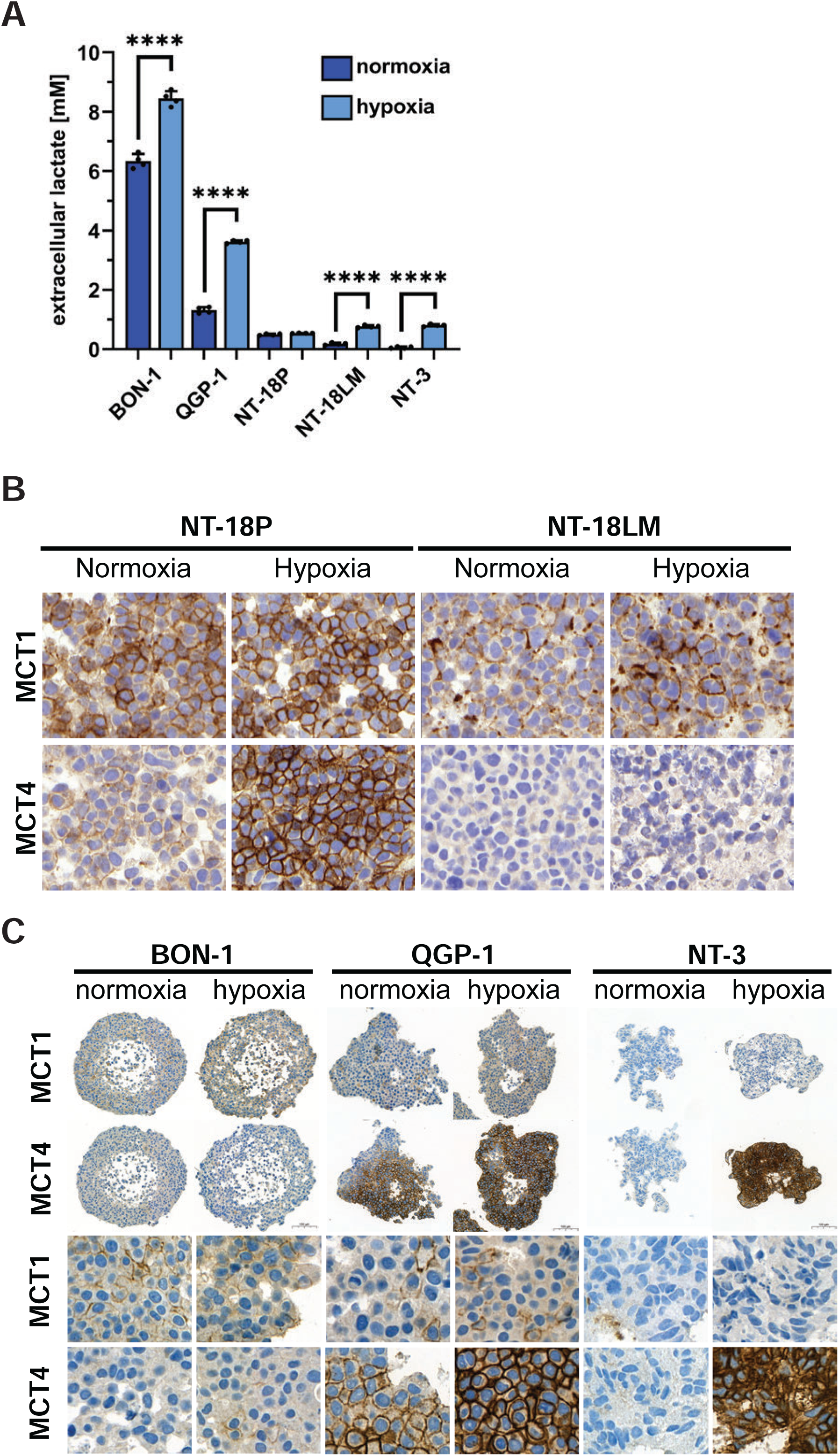
PanNET cell lines express MCT1 and MCT4 and secrete lactate. (A) Measurement of secreted lactate in conditioned media of the indicated PanNET cell lines cultured for 48h in normoxia and hypoxia (1.5% O_2_). Data are presented as mean±SD of four replicates (*p<0.05, **p<0.01, ***p<0.001, ****p<0.0001). (B) Analysis of MCT1 and MCT4 protein expression in the indicated PanNET cells and (C) spheroids cultured in normoxia and hypoxia (1.5% O_2_) for 48h by immunohistochemistry. The bottom panel in C shows sections of the spheroids at 110x magnification.

Analysis of mRNA (SFig. 3B) and protein expression (Fig. 3B) of MCT1 and MCT4 in human PanNET cell lines revealed that human PanNET cell lines express MCT1 and/or MCT4 mRNAs among other monocarboxylate transporters when cultured in 2D under normoxic conditions. Expression of HIF-1α-regulated MCT4 was significantly increased by hypoxia in all cell lines but BON-1 (SFig. 3C). Immunohistochemistry of PanNET cells demonstrated robust plasma membrane expression of MCT1 in NT-18P and NT-18LM cells grown in 2D in normoxia and hypoxia (Fig. 3B). MCT4 protein expression was detectable in NT18-P cells in normoxia, which strongly increased in hypoxia. In contrast, NT-18LM cells did not express MCT4 protein in both conditions despite considerable levels of mRNA expression (SFigs. 3B and 3C).

Under 3D culture conditions, BON-1 cells showed homogenous, low MCT1 expression throughout the spheroids in both normoxia and hypoxia (Fig. 3C). In contrast, NT-3 spheroids stained negative for MCT1, and QGP-1 spheroids displayed dispersed small clusters of MCT1-positive cells in normoxia and hypoxia (Fig. 3C). BON-1 spheroids were MCT4-negative in both oxygen conditions (Fig. 3C), whereas hypoxia led to a very strong increase in MCT4 expression in NT-3 spheroids when compared to normoxic conditions where MCT4 was undetectable. Lastly, QGP-1 spheroids displayed heterogenous MCT4 staining. MCT4 expression was strongest at the center of the spheroid and gradually diminished in intensity towards the periphery, where cells often lacked any detectable levels of MCT4. Like in NT-3 spheroids, hypoxia caused a strong increase in MCT4 expression with a homogenous staining pattern throughout QGP-1 spheroids.

In reference to the observed heterogeneity of MCT1/4 expression in both PanNET patient cohorts (Fig. 1A), human PanNET cell lines can be divided into four groups based on the conditional plasma membrane expression of MCT1 and MCT4 proteins in normoxia and hypoxia: MCT1^-^/MCT4^-^ (NT-3 cells in normoxia), MCT1^+^/MCT4^-^ (BON-1 and NT-18-LM cells in normoxia and hypoxia), MCT1^-^/MCT4^+^ (NT-3 cells in hypoxia), and MCT1^+^/MCT4^+^ (QGP-1 and NT-18P cells in normoxia and hypoxia). Thus, all patient MCT1/MCT4 expressor types are represented by cell line models. Notably, the center-to-periphery decrease in MCT4 expression in normoxic QGP-1 spheroids suggests the presence of an oxygen / HIF-1α activity gradient and metabolic heterogeneity and is reminiscent of PanNETs with heterogeneous MCT4 expression (Fig 1B).

### Inhibition of MCT1 and MCT4 impairs PanNET cell metabolism and growth

Our observation that patient tumor samples and PanNET cell lines demonstrated different subtypes based on the expression of MCT1 and MCT4 raised challenges to block lactate efflux by monotherapy. In addition, enhanced MCT4 expression was identified as an adaptive resistance mechanism to MCT1 inhibition (Curtis et al., 2017; Le Floch et al., 2011). To overcome these potential challenges, we tested syrosingopine (Fig. 4A), a clinically approved antihypertensive drug (Bartels, 1959) which has been recently repurposed for its anti-lactate efflux activity via MCT1 (IC_50_ ∼2.5μM) and MCT4 (IC_50_ ∼40nM) co-inhibition (Benjamin et al. 2018; Buyse et al. 2022). As alternative therapeutic strategies, we also tested the MCT1/MCT2 inhibitor AZD3965 (a derivative of AR-C155858) and selective MCT4 inhibitor AZD0095 (Goldberg et al., 2023) (Fig. 4A). As shown in Figure 4B, the MCT1/MCT4 dual inhibitor syrosingopine significantly reduced metabolic activity (resazurin conversion) after 48h in a dose-dependent manner in all tested human PanNET cell lines.

**Figure 4.**
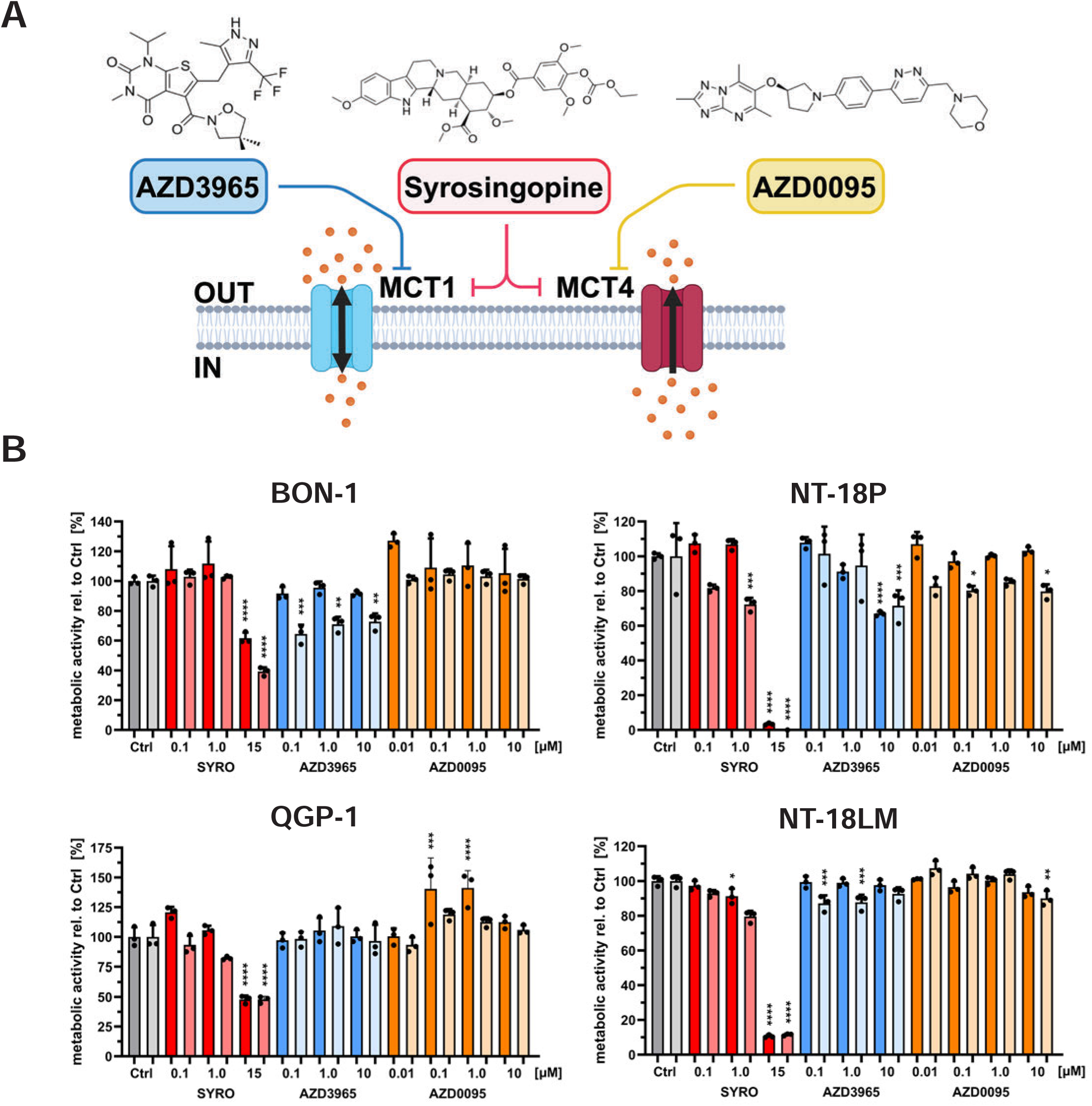
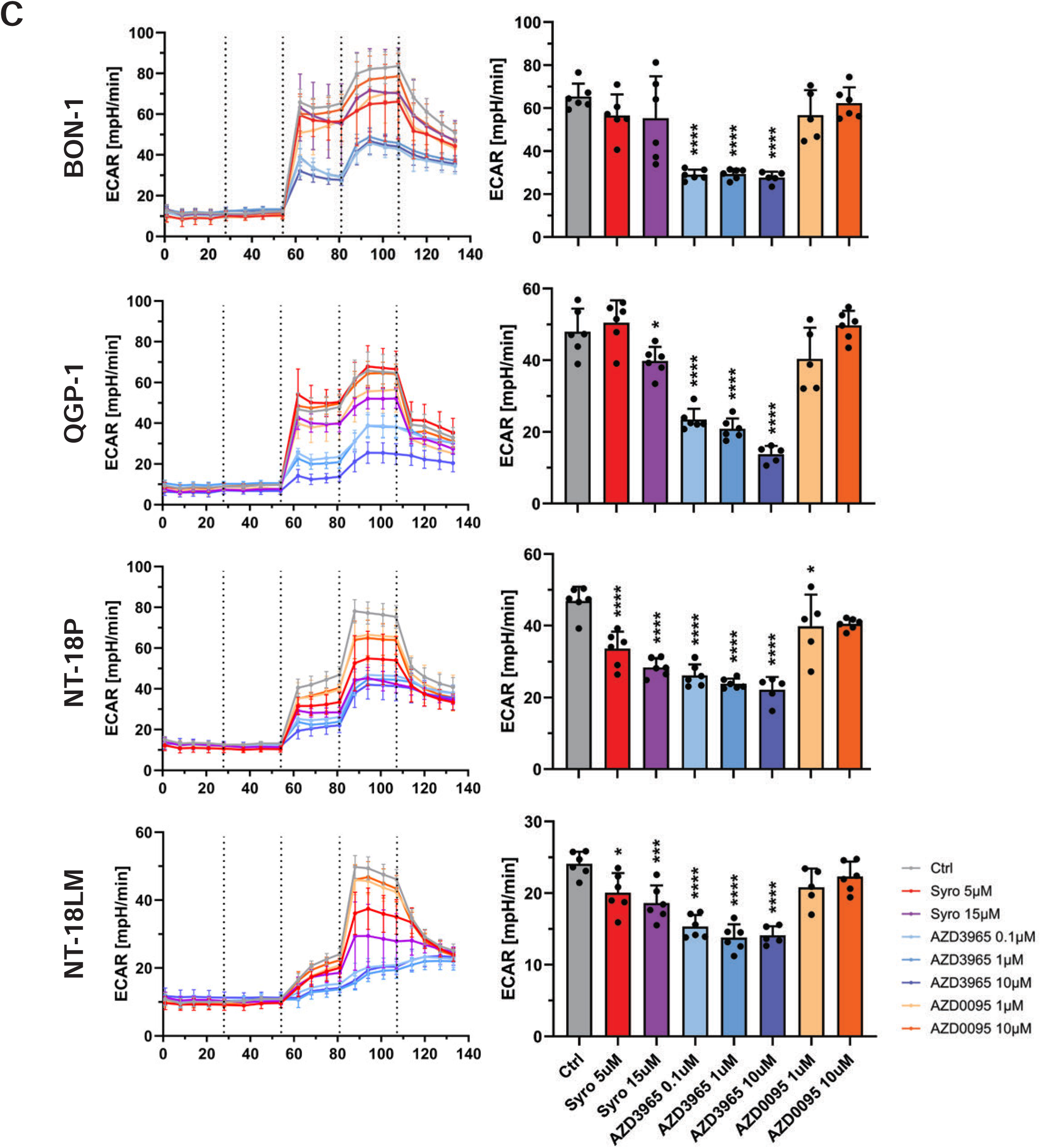
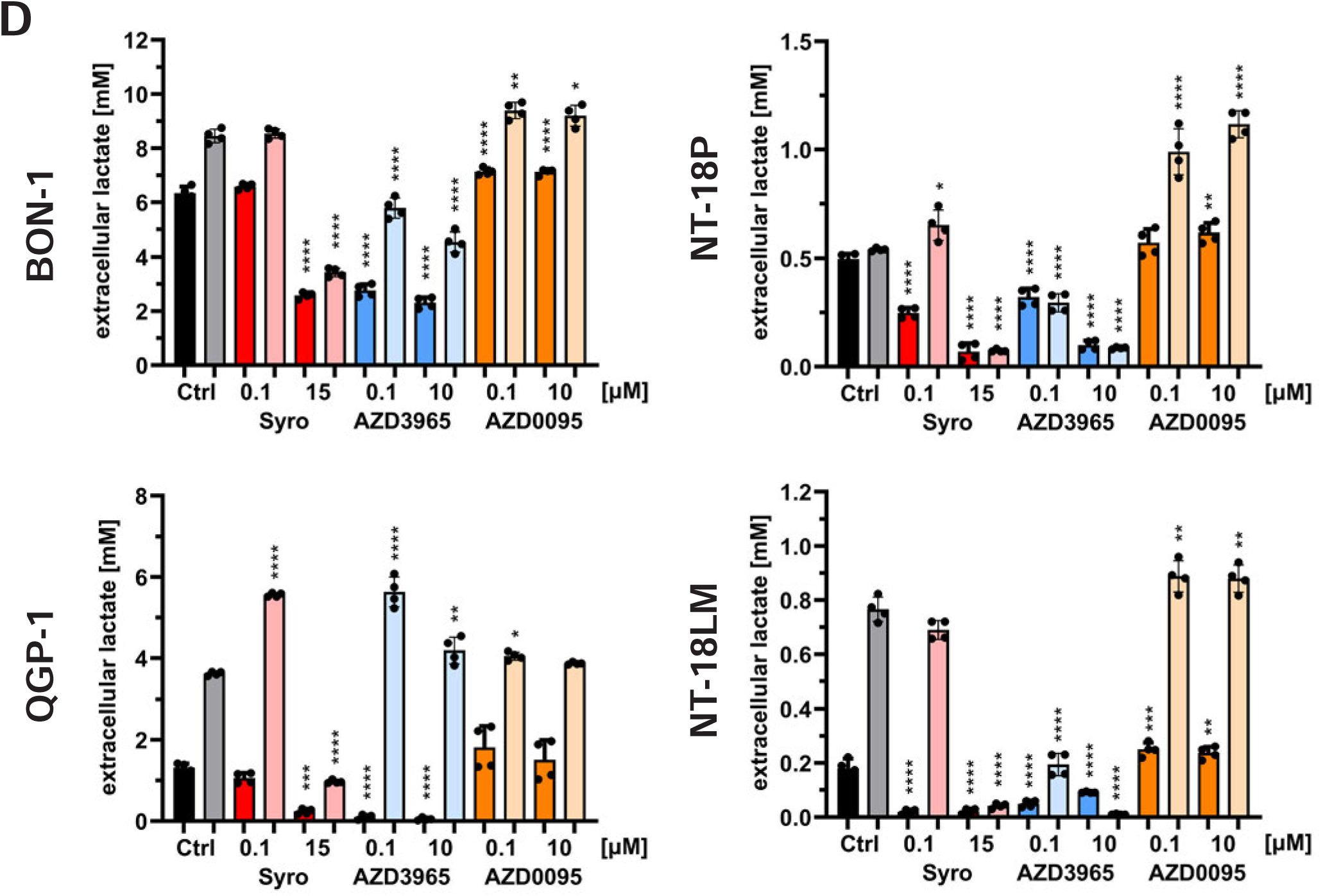
Pharmacological inhibition of MCT1 and MCT4 reduces glycolysis and lactate secretion of PanNET cells. (A) Chemical structures of the MCT4 inhibitor AZD0095, MCT1/4 inhibitor syrosingopine, and MCT1/2 inhibitor AZD3965. Graphical representation of MCT1/2/4 function, their expected lactate transport directions and inhibitor specificities. (B) After treatment of the indicated five PanNET cell lines with increasing doses of syrosingopine (SYRO), AZD3965, and AZD0095, cells were cultured in normoxia or hypoxia (1.5% O_2_) for 48h, and metabolic activity was measured by PrestoBlue assay. Data are presented as mean±SD from a representative experiment repeated three times (n=3 technical replicates). (C) Seahorse metabolic flux assay to measure extracellular acidification rate (ECAR) as a function of glycolysis and lactate secretion. The indicated PanNET cell lines were cultured for 48h in normoxia prior the start of the glycolytic stress test. Two hours before the start of the measurements, the culture media of the indicated PanNET cell lines was changed to serum- and glucose-free media and contained only glutamine to support oxidative phosphorylation (Suppl. Fig. 4B). The dashed vertical lines indicate the start of the different phases of the glycolytic stress test assay (inhibitor injection to block MCT1/2/4 activity, glucose injection to start glycolysis, ATP synthase inhibition with oligomycin for maximum glycolytic rate, glycolysis inhibition with 2-deoxyglucose). The bar charts represent the last measurement of the glucose injection phase (∼81min). Data are presented as mean±SD of 5-6 replicates. (D) Measurement of secreted lactate in conditioned media of the indicated PanNET cell lines treated for 48h with increasing doses of syrosingopine (SYRO), AZD3965, and AZD0095 in normoxia (saturated colors) and hypoxia (1.5% O_2_, pale colors). Data are presented as mean±SD of two experiments with n=4 technical replicates. Statistical significance for B-D: *p<0.05, **p<0.01, ***p<0.001, ****p<0.0001.

Consistent with HIF-1α-regulated expression of MCT4, syrosingopine’s potency was visibly stronger in hypoxia. In contrast, individual targeting of MCT1 and MCT4 with AZD3965 and AZD0095, respectively, showed only limited inhibition of metabolic activity in normoxia and hypoxia (Fig. 4B). To confirm inhibition of glycolysis and lactate export, we conducted a glycolytic stress test using the Seahorse metabolic flux analyzer (Fig. 4C). We measured the extracellular acidification rate (ECAR, mpH/min) and oxygen consumption rate (OCR, pmol/min) as functions of aerobic glycolysis (lactate secretion) and oxidative phosphorylation, respectively. In the absence of glucose, pyruvate, and lactate but presence of glutamine for oxidative phosphorylation, treatment of the PanNET cell lines BON-1, QGP-1, NT-18P and NT-18LM with the MCT inhibitors did not affect base line ECAR or OCR (t_0_-t_54_ min) in both normoxic and hypoxic conditions. This indicated that the compounds were not generally cytotoxic and that alternative MCT cargo metabolites which were potentially present in serum-free media did not fuel energy synthesis (Fig. 4C and SFig 4A). After glucose injection (t_54_ min), the MCT1/2 inhibitor AZD3965 showed the strongest inhibition of glycolysis-mediated ECAR in both normoxia and hypoxia. The dual MCT1/4 inhibitor syrosingopine caused a less severe, yet significant dose-dependent reduction in ECAR at both oxygen levels. In contrast, the selective MCT4 inhibitor AZD0095 showed the weakest anti-glycolytic activity. Notably, the inhibitory effect of AZD3965 and syrosingopine was lost in QGP-1 cells cultured in hypoxia, whereas that of AZD0095 reached significance. Given that ATP levels are maintained by balanced activities of glycolysis and oxidative phosphorylation, inhibitor-mediated glycolytic effects were inversely mirrored by changes in oxygen consumption (SFig 4A) and indicated an increased reliance of MCT1/4-compromised cells on respiration. Taken together, above results highlight that acute inhibition of MCT1/2 with AZD3965 and MCT1/4 with syrosingopine blocked lactate efflux in PanNET cells in both culture conditions, normoxia and hypoxia.

To measure lactate efflux at steady-state including adaptive compensatory effects, we measured lactate secretion of PanNET cells treated for 48h with the MCT inhibitors in normoxia and hypoxia (Fig. 4D). Notably, the longer treatment period of the lactate assay (48h, Fig. 4D) and previous metabolic assay (48h, Fig. 4A) compared to the Seahorse assay (∼1.5h, Fig. 4C) might have provided PanNET cells with sufficient time to activate compensatory mechanisms, such as expression of alternative lactate shuttles (Le Floch et al. 2011; Polański et al. 2014; Curtis et al. 2017). Thus, a longer treatment period was likely to capture the full therapeutic effect of the dual-targeting strategy as suggested by the metabolic assay (Fig. 4A). Indeed, MCT1/4 co-inhibition with syrosingopine strongly reduced lactate secretion in both normoxia and hypoxia in a dose-dependent manner in all PanNET cell lines (Fig. 4D). In normoxia, MCT1/2 inhibition with AZD3965 was similarly effective as syrosingopine. However, AZD3965’s potency was visibly reduced in BON-1 and QGP-1 cells when treated in hypoxia. The MCT4 inhibitor AZD0095 failed to reduce lactate secretion in all cell lines and conditions, except in NT-3 cells in hypoxia (data not shown). Taken together, these results demonstrate that co-inhibition of both lactate efflux systems (MCT1 and MCT4) is necessary to effectively reduce lactate secretion, in particular when MCT4 expression is expected to increase due to low oxygen levels or potential compensatory mechanisms in response to chronic MCT1 inhibition (Le Floch et al. 2011; Polański et al. 2014; Curtis et al. 2017).

Blocking lactate export causes feedback inhibition of glycolysis through intracellular lactate accumulation and cytosolic acidification (Baek et al., 2014; Draoui and Feron, 2011; Halestrap and Wilson, 2012), which in turn lowers ATP output and glycolysis-associated biosynthetic processes. The metabolic crisis induced by MCT1/4 inhibition and blocked lactate efflux is likely to induce adaptive reprogramming of key metabolic pathways. Hence, we measured functional changes in response to AZD3965 and syrosingopine by quantitative single-cell fluorescence microscopy of different metabolic sensor probes in QGP-1 cells and spheroids (Figs. 5AB). MCT1/4 inhibition with syrosingopine significantly increased lysosomal activity, glucose and fatty acid uptake, cellular lipid content (free cholesterol, neutral lipids and phospholipids, lipid droplets), and mitochondrial mass of QGP-1 cells (Fig. 5A), suggesting substantial adaptive reprogramming of energy homeostasis and nutrient supply, in particular changing lipid metabolism and mitochondrial biology. The glycolysis inhibitor 2-deoxyglucose induced very similar metabolic changes (Fig 5A), indicating that syrosingopine-mediated inhibition of lactate efflux likely drives these adaptive, compensatory responses through its anti-glycolytic effect. Consistent with above observations in the 2D cell culture system, mitochondrial mass (data not shown), lysosomal activity, ROS, and fatty acid uptake were also significantly increased in QGP-1 spheroids after treatment with syrosingopine (Fig. 5B). Notably, AZD3965 caused comparable metabolic changes in QGP-1 spheroids, in particular enhanced fatty acid uptake (Fig. 5B).

**Figure 5.**
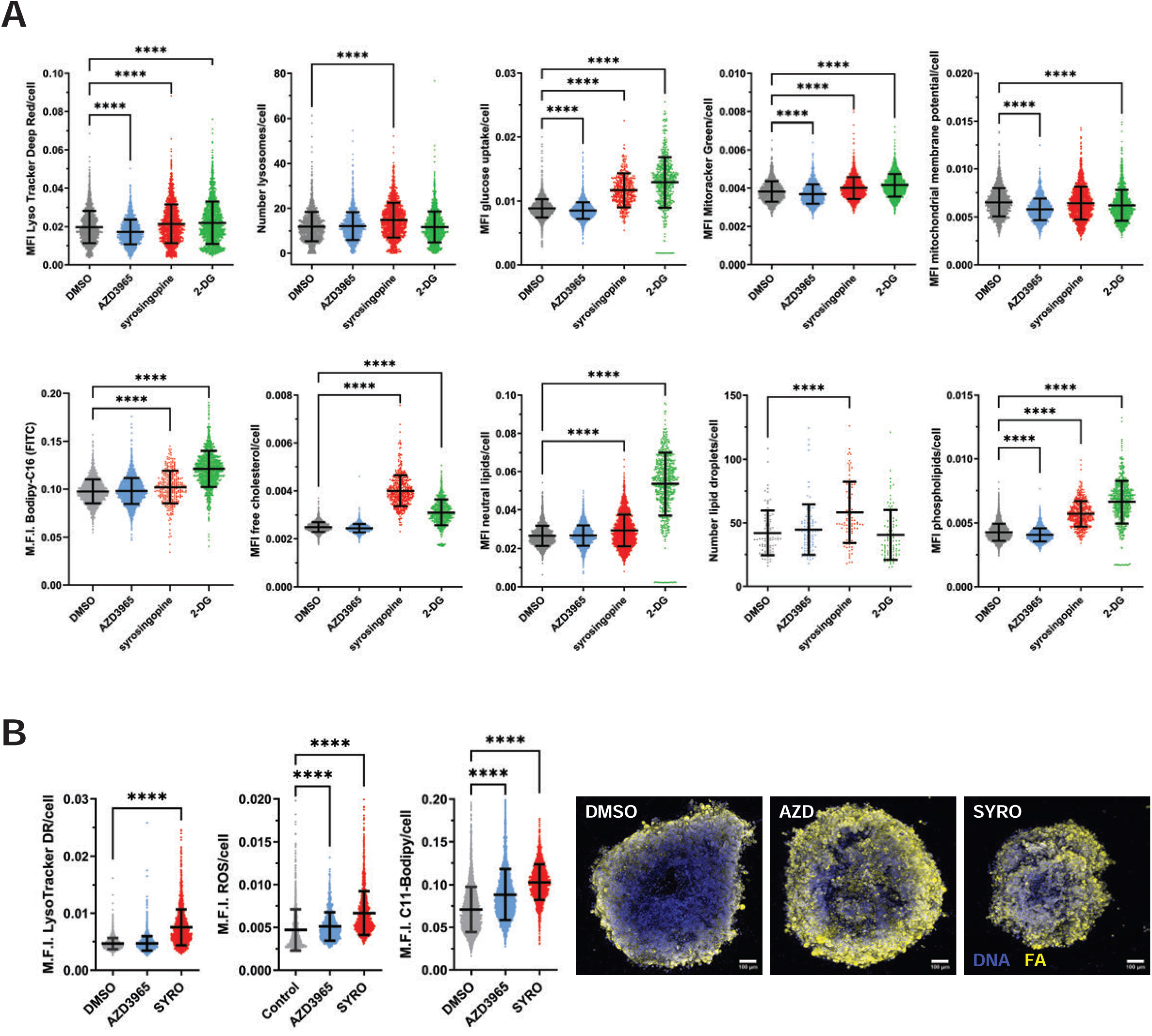
Metabolic reprogramming in response to blocked lactate efflux. (A) Functional analysis of PanNET cells treated for 48h with AZD3965 (1 μM) and syrosingopine (15 μM) using live fluorescence microscopy and quantitative single-cell analysis of lysosomal activity, glucose uptake, mitochondrial activity (mitochondrial mass and membrane potential), fatty acid uptake, and lipid content (free cholesterol, neutral lipids, lipid droplets, and phospholipids). Data are presented as mean±SD of n=3 replicates per treatment (∼2500 cells per replicate); ****p<0.0001. (B) Functional analysis of QGP-1 spheroids treated for seven days with AZD3965 (1 μM) and syrosingopine (15 μM) using live confocal fluorescence microscopy and quantitative single-cell analysis of lysosomal activity, ROS, and fatty acid (FA) uptake. Data are presented as mean±SD of n=3-4 replicates per treatment (∼3500 cells/spheroid per replicate); ****p<0.0001. Representative maximum intensity projections at 10x magnification for FA uptake (yellow) with counterstained DNA (blue) are shown.

Previous work in pancreatic adenocarcinoma, lymphoma, leukemia and breast cancer cells indicated that enhanced dependency on mitochondrial activity in response to MCT1 or MCT4 inhibition generated synthetic lethality with metformin and phenformin, both complex I inhibitors of the electron transport chain (Baek et al. 2014; Doherty et al. 2014; Hong et al. 2016; Benjamin et al. 2018). Similarly, AZD3965 and syrosingopine sensitized PanNET cells to metformin (SFig. 5), suggesting an increased dependence on mitochondrial activity when lactate efflux is compromised.

Real-time growth analysis based on time-lapse microscopy of cell confluence revealed that co-inhibition of MCT1/4 with syrosingopine significantly reduced proliferation of BON-1, QGP-1, NT-18P and NT-18LM PanNET cells (Fig. 6A and SFig. 6A). For comparison, AZD3965 (MCT1/2) caused growth inhibition in BON-1 cells but was only a weak antagonist of QGP-1 cell growth (SFig. 6A). AZD0095 (MCT4) failed to affect growth in most PanNET cell lines except BON-1 (SFig. 6A). However, combination of the MCT1/2 and MCT4 inhibitors AZD3965 and AZD0095 was more growth-suppressive than the individual compounds and showed a similar potency as syrosingopine (SFig. 6A). These results further supported the concept of dual inhibition of the MCT1/4 lactate efflux axis in PanNET. Despite the heterogeneity of MCT1 and MCT4 expression in PanNET cell lines (three groups, Fig. 3), dual inhibition of both transporters with syrosingopine successfully reduced tumor cell growth of all tested PanNET cell lines.

**Figure 6.**
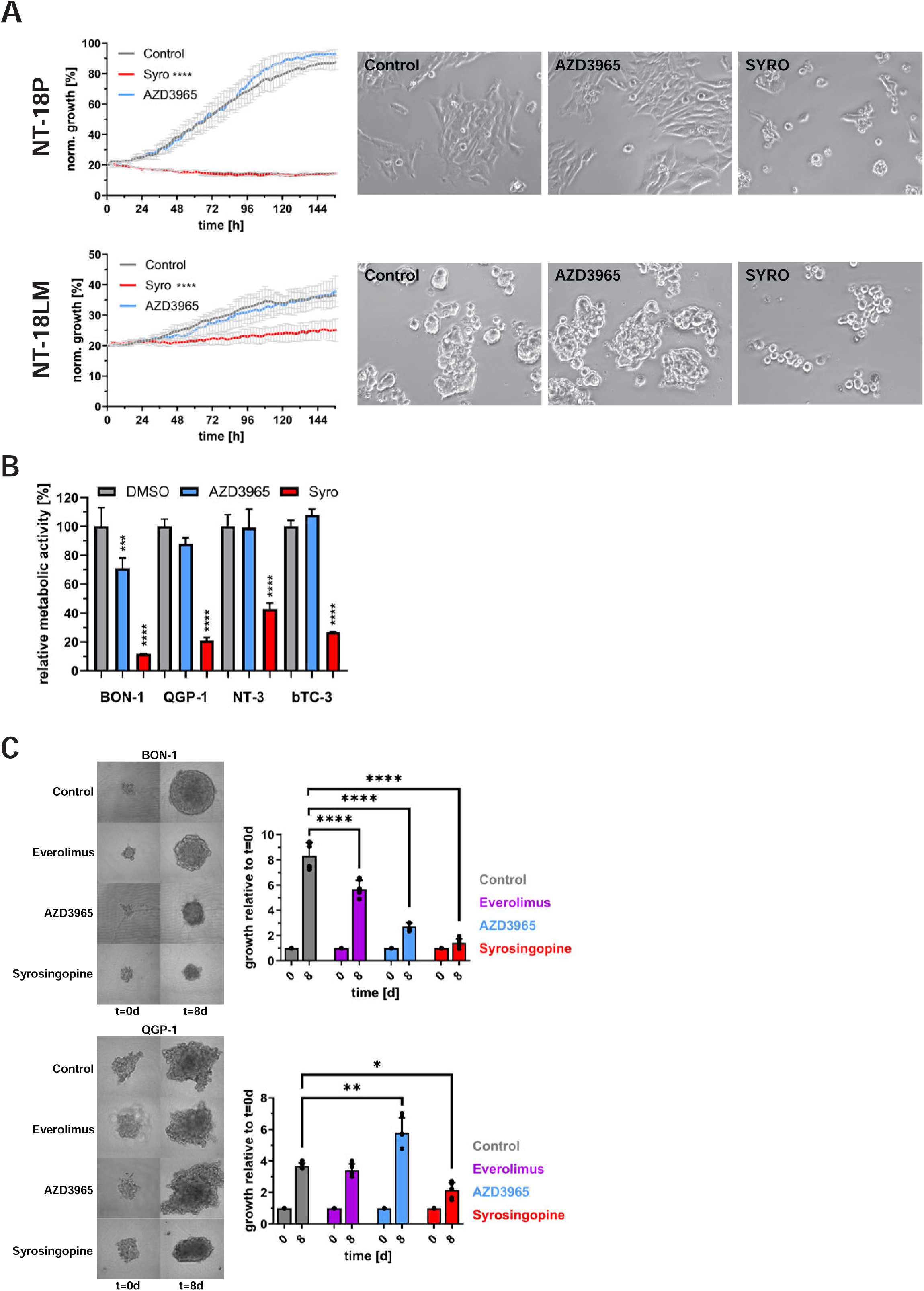
Dual targeting of MCT1 and MCT4 with syrosingopine inhibits growth of 2D and 3D PanNET models. (A) Growth of NT-18P (top panel) and NT-18LM cells (bottom panel) treated with syrosingopine (syro, 15 μM) or AZD3965 (1 μM) in normoxia was measured by label-free time-lapse microscopy of cell confluence in 2h intervals for 6.5 days. Representative micrographs of NT-18P cells and NT-18LM cells after 72h and 144h of treatment, respectively, are shown. (B) Metabolic activity of the indicated PanNET spheroids after treatment with AZD3965 (1 μM) and syrosingopine (Syro, 15 μM) for 72h was measured by PrestoBlue assay. (C) Relative growth of BON-1 and QGP-1 spheroids treated with AZD3965 and syrosingopine in normoxia was measured by label-free time-lapse microscopy of spheroid area for 8 days. mTOR1 inhibitor everolimus, a clinically approved therapy against metastatic PanNET, was used as a control. Representative micrographs of the spheroids at the specified time points of the analysis are shown. Data from a representative experiment of at least two repeats are presented as mean±SD of n=3 wells. (*p<0.05, **p<0.01, ***p<0.001, ****p<0.0001).

### Co-inhibition of MCT1 and MCT4 blocks PanNET spheroid and tumoroid growth

Compared to 2D cell culture, cells within 3D cell culture models, e.g., spheroids, have increased cell-cell contacts and experience regional differences in oxygen and nutrient supply as well as treatment exposure depending on how deep they are positioned within the spheroid. Thus, treatment of PanNET spheroids might provide valuable insights into the efficacy of MCT1/4 inhibition in the context of regional heterogeneity (core versus periphery of the spheroid, Figs. 3BC and SFig. 6E) of a more complex model. AZD3965 decreased metabolic activity specifically in BON-1 spheroids (Fig. 6B), whereas syrosingopine showed a strong inhibitory effect in all tested spheroids, including both human (BON-1, QGP-1, NT-3) and mouse PanNET spheroids (betaTC-3). Consistent with increased lactate synthesis and efflux in oxygen-deprived conditions, syrosingopine’s potency was even higher in spheroids which were cultured in hypoxia (SFig. 6B). Time-lapse microscopy showed that the growth of BON-1 and QGP-1 spheroids was significantly reduced by the MCT1/4 inhibitor syrosingopine (Fig. 6C and SFigs. 6C and 6E). Inhibition of MCT1/2 with AZD3965 showed cell-line specific effects; suppression of growth in BON-1 and, surprisingly, stimulation of growth in QGP-1 spheroids (Fig. 6C). Notably, AZD0095 showed very little inhibitory activity (data not shown). Assessment of cell death revealed a modest but significant increase in cell death in BON-1 spheroids treated with AZD3965 and syrosingopine for four and seven days (SFigs. 6D and 6E), whereas the viability of QGP-1, NT-18P and NT-18LM spheroids was not significantly affected by either inhibitor (data not shown), suggesting that inhibition of lactate efflux was cytostatic rather than cytotoxic. Taken together, MCT1/4 dual antagonist syrosingopine showed robust inhibitory activity in PanNET 3D cell culture models.

Next, we tested syrosingopine in our PanNET patient-derived tumoroid (PDT) model (April-Monn et al. 2020; April-Monn et al. 2024) using cryopreserved tumor material from eight patients. Based on the the expression of MCT1 and/or MCT4 in cells used for seeding of the tumoroids, the eight PDTs can be divided into three groups: MCT1^+^/MCT4^-^ (n=2), MCT1^+^/MCT4^+^(n=4), MCT1^-^/MCT4^+^ (n=2). As shown in Figure 7, dual inhibition of MCT1 and MCT4 by syrosingopine significantly reduced metabolic activity in all tested PDTs. Syrosingopine’s efficacy was nearly as potent as that of multi-targeted receptor tyrosine kinase inhibitor sunitinib and ERK1/2 inhibitor SCH772984 (SFig. 7A). In support of our dual targeting strategy against MCT1 and MCT4, MCT1/2 inhibitor AZD3965 failed to show inhibitory activity in PDTs which expressed both MCT1 and MCT4 (SFig. 7A). For a more detailed phenotypic analysis of treatment response and adaptive changes within the PDTs, we assessed at treatment endpoint multiple live parameters, such as tumoroid size, cell death, and mitochondrial and lysosomal activities by confocal fluorescence microscopy and automated image analysis (Table 5 and SFig 7B). Consistent with our cell line data, syrosingopine exerted mostly a cytostatic effect, with only one out of eight PDTs showing significantly increased cell death compared to vehicle control (DMSO). The change in mitochondrial membrane potential (MMP) was heterogenous but seemed to be exclusively inhibited in MCT1/MCT4 double-positive PDTs. Like QGP-1 and BON-1 cells (Fig. 5), almost all PDTs responded to syrosingopine with significantly increased lysosomal activity, suggesting reprogramming of nutrient supply and energy homeostasis. Taken together, dual targeting of MCT1 and MCT4 with syrosingopine showed promising inhibitory activity in patient-derived PanNET tumoroids.

**Figure 7.**
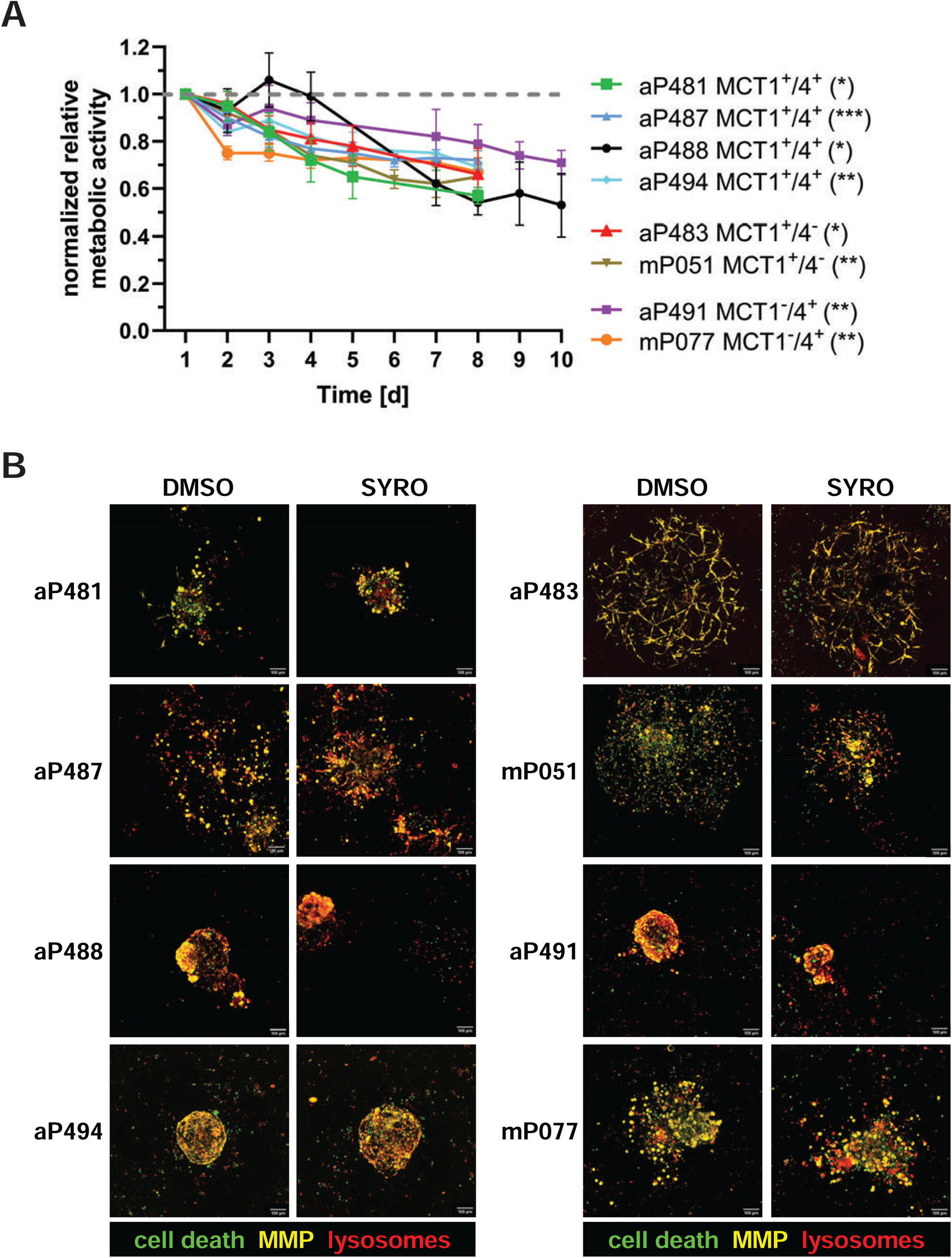
Dual targeting of MCT1 and MCT4 with syrosingopine inhibits metabolic activity of patient-derived tumoroids. (A) Real-time analysis of PDT viability based on the relative metabolic activity (reductive power) of syrosingopine-treated (15 μM). The PDTs are divided into three subtypes based on the expression of MCT1 and MCT4. The dashed line indicates the normalized, relative metabolic activity of the corresponding PDTs treated with vehicle control (DMSO) for each time point. Data are presented as mean±SD of n=3-4 replicates per treatment; *p<0.05, **p<0.01, ***p<0.001. (B) Representative confocal fluorescent images (maximum intensity projections) of control (DMSO) and syrosingopine-treated (SYRO) patient-derived tumoroids after live staining of DNA (blue), cell death (green), mitochondrial membrane potential (yellow), and lysosomal activity (red) at the endpoint of the 3D cell culture experiment (8-10 days).

**Table 5.** Summary of syrosingopine-induced changes in PDTs.

|  |  |  | syrosingopine |  |  |  |
| --- | --- | --- | --- | --- | --- | --- |
| patient | target status |  | reduced size PDT | increased cell death | changed MMP | increased lysosomal activity |
| aP481 | MCT1 | MCT4 |  |  | (-) **** | **** |
| aP487 | MCT1 | MCT4 |  |  | (-) **** | **** |
| aP488 | MCT1 | MCT4 | **** | * | (-) **** |  |
| aP494 | MCT1 | MCT4 |  |  | (-) **** | **** |
| aP483 | MCT1 | - |  |  |  | **** |
| mP051 | MCT1 | - |  |  | (+) **** | **** |
| aP491 | - | MCT4 | ** |  | (+) **** | **** |
| mP077 | - | MCT4 | *** |  |  | **** |

## Discussion

Enhanced expression of MCT1 and MCT4 and their association with adverse prognostic parameters has been previously reported in multiple types of cancer. Our analysis of two independent patient cohorts demonstrated that PanNETs show frequent expression of MCT1 or MCT4 and in some cases co-expression of both lactate shuttles. Overlapping regional co-expression suggests a cooperative function of MCT1 and MCT4 in lactate efflux, whereas mutually exclusive expression of both transporters in adjacent tumor regions, including the stroma, has been previously associated with metabolic symbiosis through lactate exchange between MCT4-positive glycolytic regions (Warburg effect or hypoxia) and MCT1-positive oxidative regions, (Allen et al., 2016; Curry et al., 2013; Mikkilineni et al., 2017). However, without the technically challenging confirmation of the lactate transport direction *in vivo* or verification of co-expression of markers of glycolysis (e.g., LDHA) and lactate oxidation (e.g., LDHB), respectively, it is speculative to assign MCT1-positive tumor regions a specific metabolic role, i.e., glycolytic (lactate efflux) or oxidative (lactate uptake). Since protons are co-secreted with lactate and acidify the extracellular space, maintenance of the acid-base balance in the extracellular space through enzymes, such as CA9, is critical for glycolytic and hypoxic tumors. Indeed, we observed good correlation of co-expression of CA9 and MCT4, indicating the potential presence of regional hypoxia and anaerobic glycolysis. The possibility of regional hypoxia was further supported by a significantly lower MVD, a marker of aggressive PanNETs (Battistella et al., 2022; Couvelard et al., 2005), in tumors with heterogenous MCT4 expression. This was in stark contrast to the frequent occurrence of homogeneous MCT4 expression in PanNET cores with high MVD, suggesting pseudohypoxia. The above observed dichotomies strongly suggest that mutually exclusive expression of MCT1 and MCT4 and homogeneous and heterogeneous MCT1/MCT4 co-expression patterns might define different metabolic subtypes (oxidative, glycolytic, hypoxic, and pseudohypoxic) within PanNETs; and as indicated by their correlation with multiple clinico-pathologically parameters of aggressive disease, that they are of clinical relevance. Thus, metabolic heterogeneity, plasticity and functional redundancy are challenges that a therapeutic strategy against lactate transport faces in PanNET.

We addressed some of these challenges in our *in vitro* cell culture studies in 2D and 3D with PanNET cell lines and patient-derived tumoroids. While all four clinically observed MCT1/4 expressor types were represented by at least one PanNET cell line when cultured in normoxic or hypoxic conditions, the direction of lactate transport was functionally less complex than expected, i.e., unidirectional in all examined conditions. AZD3965-mediated loss of MCT1 function reduced extracellular acidification and lactate secretion in all four tested PanNET cell lines, assigning MCT1 a critical glycolytic role. This functional dependency was lost or reduced in hypoxia, where higher lactate efflux was mediated through MCT4. These observations indicate a unidirectional and cooperative function of MCT1 and MCT4 in mediating lactate efflux in PanNET cells *in vitro*. Similarly, MCT4 expression has been shown to confer resistance to MCT1 inhibition in various cancer cell lines, implicating MCT1 in lactate efflux (Le Floch et al. 2011; Polański et al. 2014; Curtis et al. 2017). Consistent with this functional redundancy, co-inhibition of MCT1 and MCT4 with syrosingopine was more efficacious in reducing metabolic activity and growth of PanNET cells, spheroids and patient-derived tumoroids than inhibition of MCT1 and MCT4 individually with either AZD3965 or AZD0095. In particular syrosingopine’s potency in more complex 3D cell culture models of PanNET with visible heterogeneity in MCT1/4 protein expression (Fig. 3C) and metabolic features (Fig. 5B, SFig. 6E, and Fig. 7B) are promising indications that syrosingopine might be effective against all three clinical expressor types (MCT1^+^/MCT4^-^, MCT1^-^/MCT4^+^ and MCT1^+^/MCT4^+^) *in vivo* as was observed *in vitro* in the PDT model.

Metabolic plasticity is a hallmark of cancer and limits the efficacy of many inhibitors directed against metabolic pathways. Our work revealed adaptive metabolic reprogramming of multiple pathways (glucose and fatty acid uptake, lipid metabolism) and organelles (mitochondria, lysosomes and lipid droplets) in response to the anti-glycolytic effects of blocked lactate efflux. Adaptive resistance mechanisms, such as upregulation of alternative lactate shuttles (Le Floch et al. 2011; Polański et al. 2014; Curtis et al. 2017), nutrient scavenging through macropinocytosis, enhanced autophagy or rebalancing of energy homeostasis through increased oxidative phosphorylation (OxPhos) of mitochondrial fuels, such as pyruvate and glutamine have been previously shown to partially rescue cancer cells from MCT1/4 inhibition (Baek et al. 2014). Consistent with enhanced mitochondrial activity, mitochondrial mass and ROS were increased in pancreatic adenocarcinoma cells in response to MCT4 inhibition (Baek et al., 2014). However, such stress-induced reprogramming can generate metabolic vulnerabilities, as previously demonstrated by the lethal effects of OxPhos inhibitors metformin, phenformin or rotenone when combined with MCT1 or MCT4 inhibition in pancreatic adenocarcinoma, lymphoma, leukemia and breast cancer cells (Baek et al., 2014; Benjamin et al., 2018; Doherty et al., 2014; Hong et al., 2016). Likewise, AZD3965 and syrosingopine sensitized PanNET cells to metformin, indicating that PanNET cells become critically dependent on mitochondrial ATP generation when lactate efflux is impaired.

Impairing lactate efflux might have additional therapeutic benefits for PanNET patients. Since lactate is a known immunosuppressive metabolite (Wang et al., 2021), suppression of lactate efflux via MCT1/4 co-inhibition might increase the immune cell activity in PanNET. MCT1/4 dual targeting with syrosingopine might be particularly effective *in vivo* when combined with current PanNET therapy sunitinib, which targets multiple receptor tyrosine kinases, including VEGF, and blocks angiogenesis. The benefit of inhibited angiogenesis would be twofold: Firstly, it causes hypoxia in the tumor and enhances glycolysis and therefore the expression of MCT4 (Allen et al., 2016) and dependence on lactate efflux, Secondly, it preemptively counters adaptive metabolic reprogramming of syrosingopine-treated tumor cells (Fig. 5) by restricting access to alternative fuels such as glutamine or fatty acids for mitochondrial OxPhos, thereby increasing energy stress and cytotoxicity.

A limitation of our study is that we could not test syrosingopine in an appropriate *in vivo* setting since a mouse model for non-functioning PanNET has not been established yet. In addition, there is currently a scarcity of human PanNET cell lines available. We addressed these challenges by our investigations in 2D and 3D cell culture models with demonstrated metabolic heterogeneity. More importantly, our tumoroid model using patient-derived tumor cells was extremely valuable for testing our dual targeting concept against lactate efflux in a clinically relevant 3D model of PanNET. Unfortunately, our cryo-sample collection does not contain material from tumors which are double-negative for MCT1/MCT4 expression. Thus, we were unable to compare the specificity of syrosingopine to the three MCT1/MCT4 expressor types. Furthermore, despite our TMAs incorporating PanNET patient samples of two academic centers, the case numbers were too small for some comparisons to reach statistical significance.

In conclusion, our study demonstrated that PanNET is metabolically a very heterogeneous neoplasm. We classified four metabolic subtypes based on MCT1 and MCT4 expression, but additional subtypes are likely to exist within the largest group which lacked expression of both lactate shuttles. Our 2D and advanced 3D cell culture studies showed that dual inhibition of MCT1 and MCT4 is a promising therapeutic strategy against several metabolic PanNET subtypes.

## Acknowledgments

This work was funded by a Swiss National Science Foundation grant 310030-188639 to A. Perren. Microscopy was performed on equipment supported by the Microscopy Imaging Center Bern. KB was supported by a Swiss National Science Foundation Postdoctoral Fellowship grant (award number P500PM_217647 / 1).

## Conflict of Interest

None.

---

**Supplementary Table 1.** Primer sequences.

**Supplementary Table 2.** Summary of inhibitor-mediated changes in PDTs.

**Supplementary Figure 1.** MCT1/4 protein co-expression. Representative tissue cores of four MCT1 and MCT4 double-positive PanNETs demonstrating strong regional co-expression (tissue cores on the right) as well as regions with exclusive MCT1 or MCT4 expression in subsequent slides from the same tissue core (tissue cores on the left).

**Supplementary Figure 2.** Violin plots of Ki67 index, microvessel density (MVD), and tumor size of MCT1-negative, -heterogeneous positive and -homogeneous positive PanNETs of the Bern (top panels) and Milano cohorts (bottom panels). MCT1-expressors tend to be larger (non-significant) and demonstrate higher proliferation (non-significant). Heterogeneous expression associates with lower MVD (non-significant). *TC*: Tumor Center; *TF*: Tumor Front.

**Supplementary Figure 3.** (A) PrestoBlue assay to measure metabolic activity as a function of cellular reductive power of the indicated PanNET cell lines cultured for 48h in normoxia (blue bars) and hypoxia (1.5% O_2_, red bars). Data are presented as mean±SD of three independent experiments (n=3 technical replicates). (B) mRNA expression confirmed lactate shuttle genes (MCT1-4) and HIF1α-regulated genes in the indicated PanNET cell lines cultured in normoxia and hypoxia for 48h were determined my RNAseq. Missing bars indicate undetectable expression. (C) Quantitative analysis of MCT4 mRNA expression by qRT-PCR of the indicated cell lines cultured in normoxia and hypoxia (1.5% O_2_) for 48h. Data are presented as mean±SD of two independent experiments (n=3 technical replicates, *p<0.05, **p<0.01, ***p<0.001, ****p<0.0001).

**Supplementary Figure 4.** Seahorse metabolic flux assay to measure (A) extracellular acidification rate (ECAR) as a function of glycolysis and lactate secretion in hypoxia and (B) oxidative consumption rate (OCR) as a function of oxidative phosphorylation in normoxia and hypoxia. The indicated PanNET cell lines were cultured for 48h in normoxia or hypoxia (1.5% O_2_) prior the start of the glycolytic stress test. Two hours before the start of the measurements, the culture media of the indicated PanNET cell lines was changed to serum- and glucose-free media and contained only glutamine to support oxidative phosphorylation. The dashed vertical lines indicate the start of the different phases of the glycolytic stress test assay (inhibitor injection to block MCT1/2/4 activity, glucose injection to start glycolysis, ATP synthase inhibition with oligomycin for maximum glycolytic rate, glycolysis inhibition with 2-deoxyglucose) which was also carried out in normoxia and hypoxia (1.5% O_2_). The bar charts represent the last measurement of the glucose injection phase (∼81min). Data are presented as mean±SD of 5-6 replicates. *p<0.05, **p<0.01, ***p<0.001, ****p<0.0001.

**Supplementary Figure 5.** (A) Growth of BON-1 (top panel) and QGP-1 cells (bottom panel) treated with AZD3965 (1 μM) and AZD0095 (1 μM) individually or in combination was measured by label-free time-lapse microscopy of cell confluence in 2h intervals for 5 days. Data are presented as mean±SD of n=3 wells.

**Supplementary Figure 6.** (A) Growth of BON-1 (left panel) and QGP-1 cells (right panel) treated with increasing concentrations of AZD3965, AZD0095 and syrosingopine in normoxia was measured by label-free time-lapse microscopy of cell confluence in 2h intervals for 5 days. Data are presented as mean±SD of n=3 wells. (B) Metabolic activity of the indicated PanNET spheroids after treatment with syrosingopine (Syro, 15 μM) and glycolysis inhibitor 2-deoxyglucose (2-DG, 15 μM) for 72h in normoxia and hypoxia (1.5% O_2_) was measured by PrestoBlue assay. (C) Relative growth of BON-1 spheroids treated with AZD3965 (1 μM) and syrosingopine (15 μM) in normoxia was measured by label-free time-lapse microscopy of spheroid area in 8h intervals for 7 days. mTOR1 inhibitor everolimus, a clinically approved therapy against metastatic PanNET, was used as a control. (D) Relative cell death in BON-1 spheroids treated with AZD0095 (1 μM), AZD3965 (1 μM), and syrosingopine (Syro, 5 μM and 15 μM) for 24h and 96h was measured by live confocal fluorescence microscopy and quantitative single-cell analysis of Hoechst 33342- (blue, total number of nuclei) and SytoxGreen-stained spheroids (green, nuclei of dead cells) with CellProfiler software. Representative maximum intensity projections of spheroids treated with vehicle control (DMSO), AZD3965 (1 μM) and syrosingopine (SYRO, 15 μM) are shown. (E) Functional analysis of BON-1 spheroids treated for seven days with AZD3965 (1 μM) and syrosingopine (15 μM) using live confocal fluorescence microscopy and quantitative analysis of spheroid area, relative cell death and mitochondrial membrane potential (MMP). Representative maximum intensity projections at 10x magnification for mitochondrial membrane potential (MMP, yellow) and dead cells (green) are shown. Data are presented as mean±SD of 3 replicates per treatment, ns=not significant, *p<0.05, **p<0.01, ***p<0.001, ****p<0.0001.

**Supplementary Figure 7.** (A) Real-time analysis of PDT viability based on the relative metabolic activity (reductive power) of SCH772984 (SCH, 2 μM, aP488, aP491), sunitinib (SUN, 5 μM, aP481, aP487, aP494, aP483, mP051, mP077) and AZD3965-treated (1 μM, bottom panel). The PDTs are divided into three subtypes based on the expression of MCT1 and MCT4. The dashed line indicates the normalized, relative metabolic activity of the corresponding PDTs treated with vehicle control (DMSO) for each time point. Data are presented as mean±SD of n=3-4 replicates per treatment; ns=not significant, *p<0.05, **p<0.01, ***p<0.001. (B) Representative confocal fluorescent images (maximum intensity projections) of sunitinib (SUN) and SCH772984-treated (SCH) patient-derived tumoroids after live staining of DNA (blue), cell death (green), mitochondrial membrane potential (yellow), and lysosomal activity (red) at the endpoint of the 3D cell culture experiment (8-10 days).

## Notes

### Competing Interest Statement

The authors have declared no competing interest.

