## Supplementary Table S1 for "Dual inhibition of lactate transporters MCT1 and MCT4 in pancreatic neuroendocrine tumors targets metabolic heterogeneity and functional redundancy"

Supplementary Table 1: Primer sequences

| Gene |  | 5’-Sequence-3’ |
| --- | --- | --- |
| RPL32 | Forward | GCACCAGTCAGACCGATATG |
|  | Reverse | ACTGGGCAGCATGTGCTTTG |
| MCT4 (SLC16A3) | Forward | ACAGGTGAGGCGGAACCAA |
|  | Reverse | CAGTGATGACGAAACAGCCGA |
