## Supplementary Table S2 for "Dual inhibition of lactate transporters MCT1 and MCT4 in pancreatic neuroendocrine tumors targets metabolic heterogeneity and functional redundancy"

| sunitinib / SCH772984 (x) | | | | | | |
|---|---|---|---|---|---|---|
| **patient** | **target status** | | **reduced size PDT** | **increased cell death** | **changed MMP** | **increased lysosomal activity** |
| aP481 | MCT1 | MCT4 | **** | | (-) **** | **** |
| aP487 | MCT1 | MCT4 | | *** | (-) **** | **** |
| aP488 x | MCT1 | MCT4 | **** | | | **** |
| aP494 | MCT1 | MCT4 | **** | *** | (-) **** | **** |
| aP483 | MCT1 | - | **** | ** | (-) **** | **** |
| mP051 | MCT1 | - | * | * | (+) **** | **** |
| aP491 x | - | MCT4 | ** | | (-) **** | *** |
| mP077 | - | MCT4 | **** | **** | (+) **** | **** |
