## Supplementary Figures S1 to S7 for "Dual inhibition of lactate transporters MCT1 and MCT4 in pancreatic neuroendocrine tumors targets metabolic heterogeneity and functional redundancy"

### Suppl Figure 1

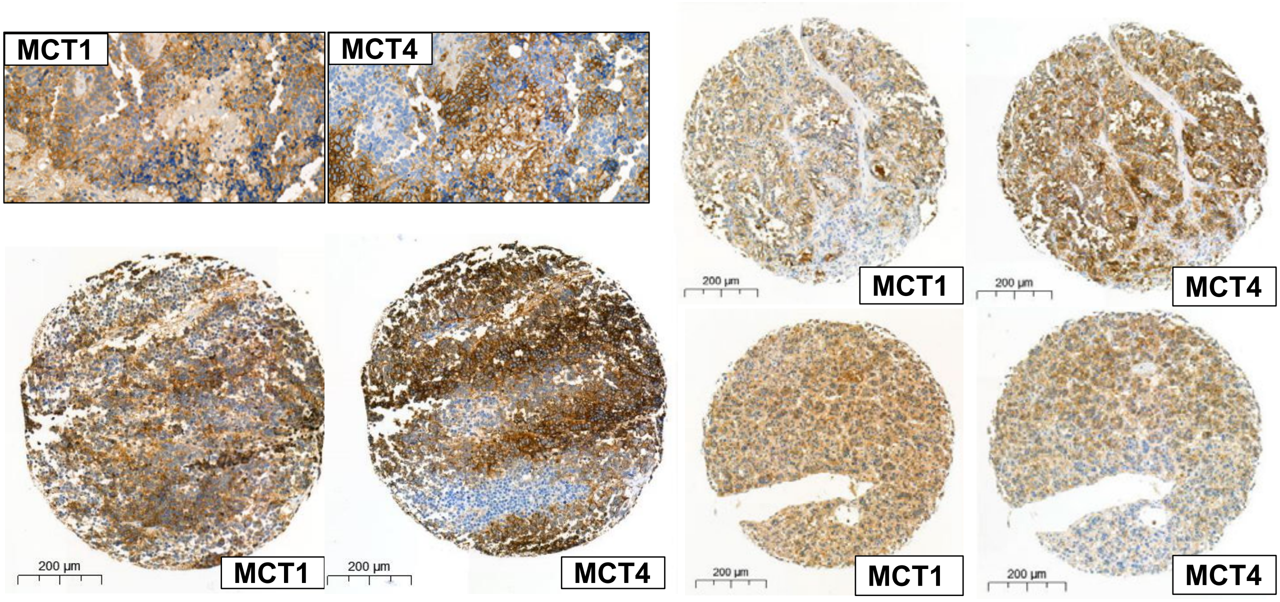

### Suppl Figure 2

A

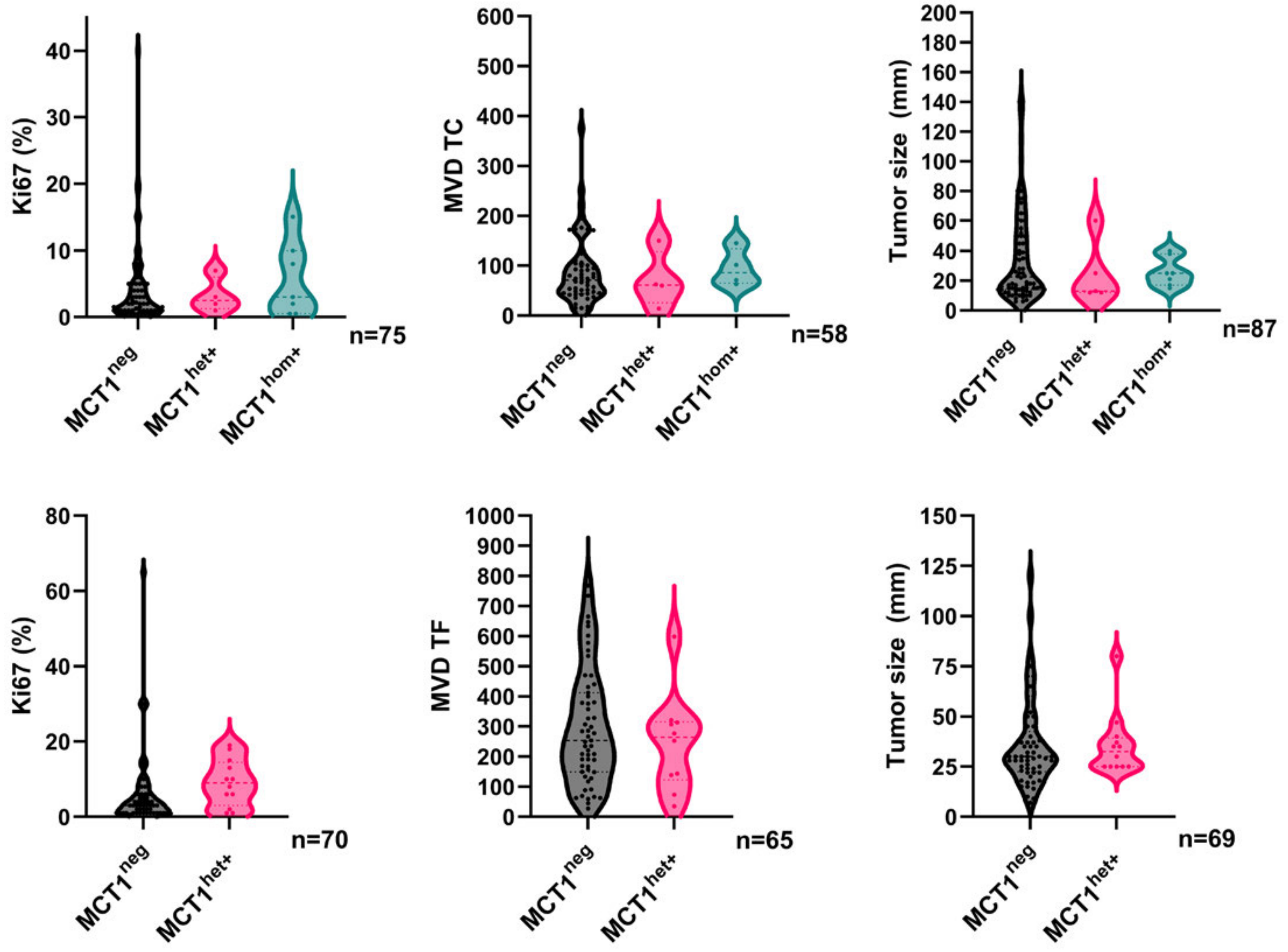

### Suppl Figure 3

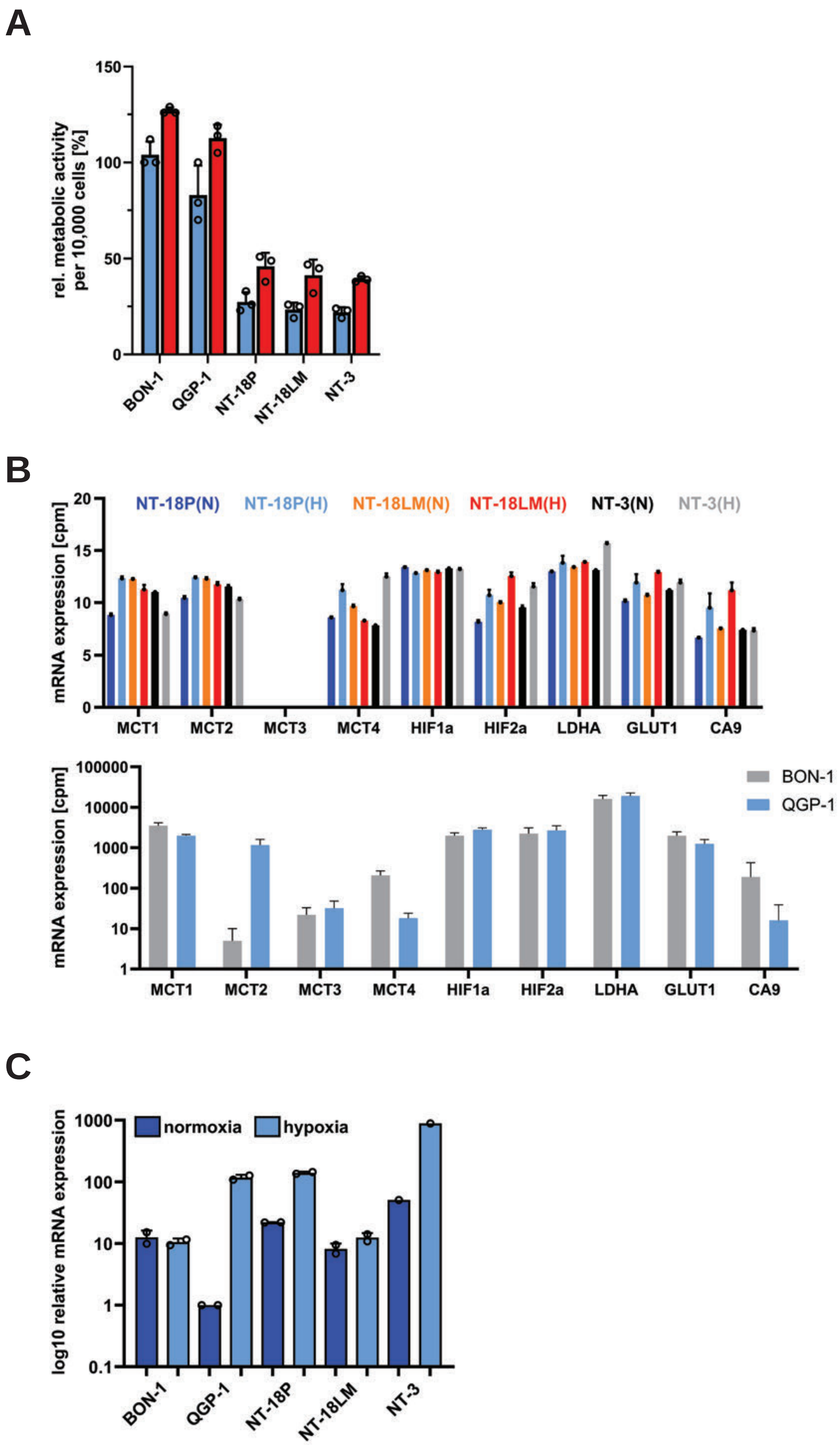

**Suppl. Figure 4**

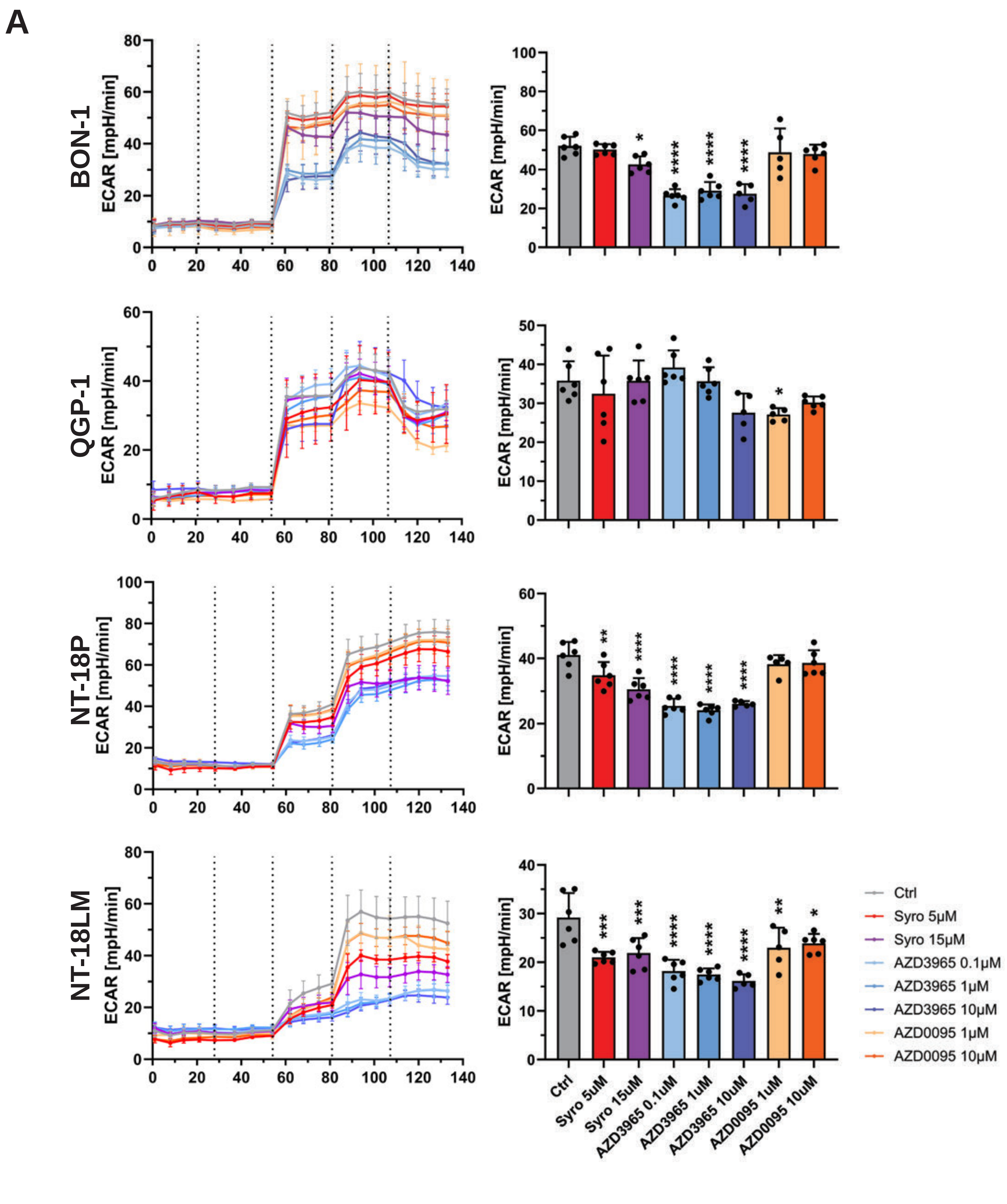

#### Suppl Figure 4

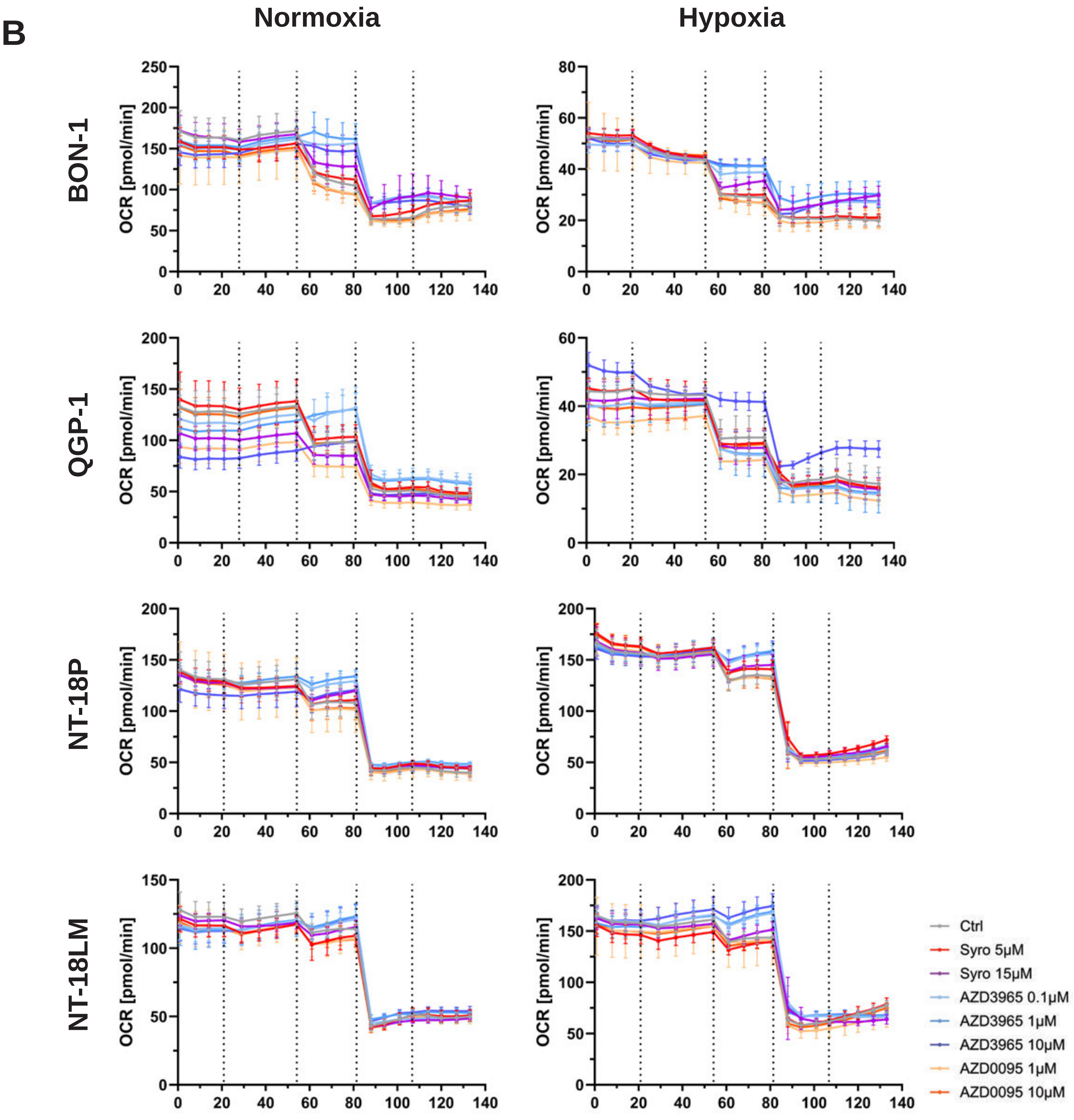

### Suppl Figure 5

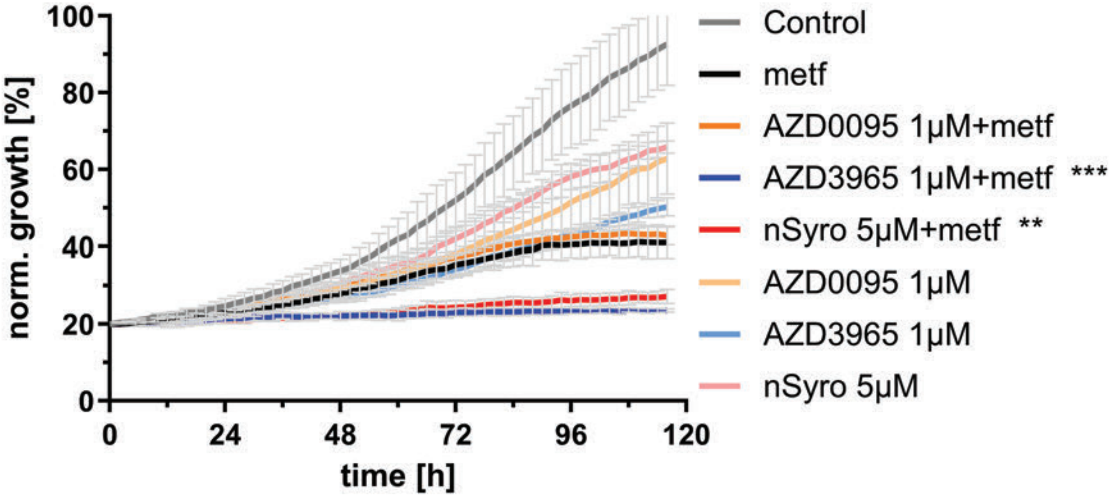


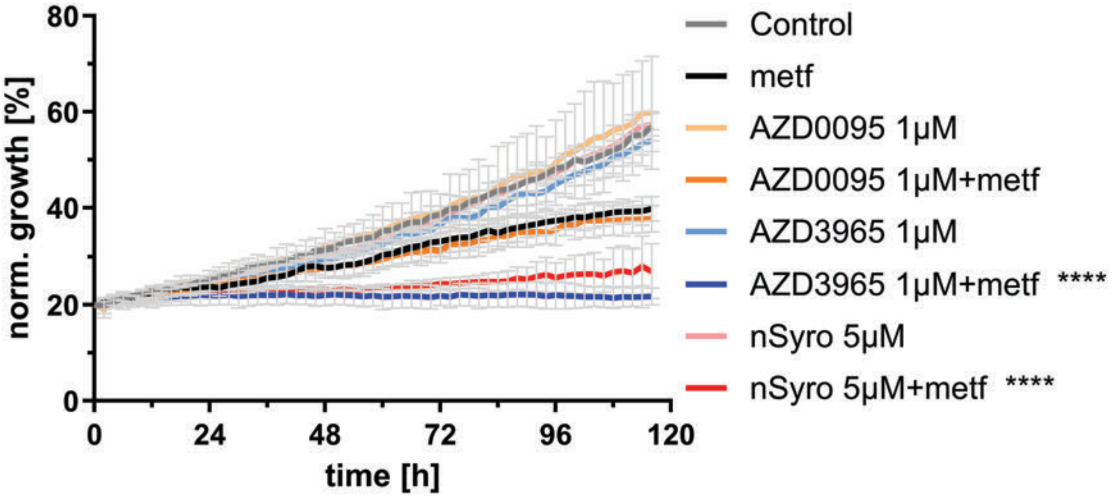

### Suppl Figure 6

**A**

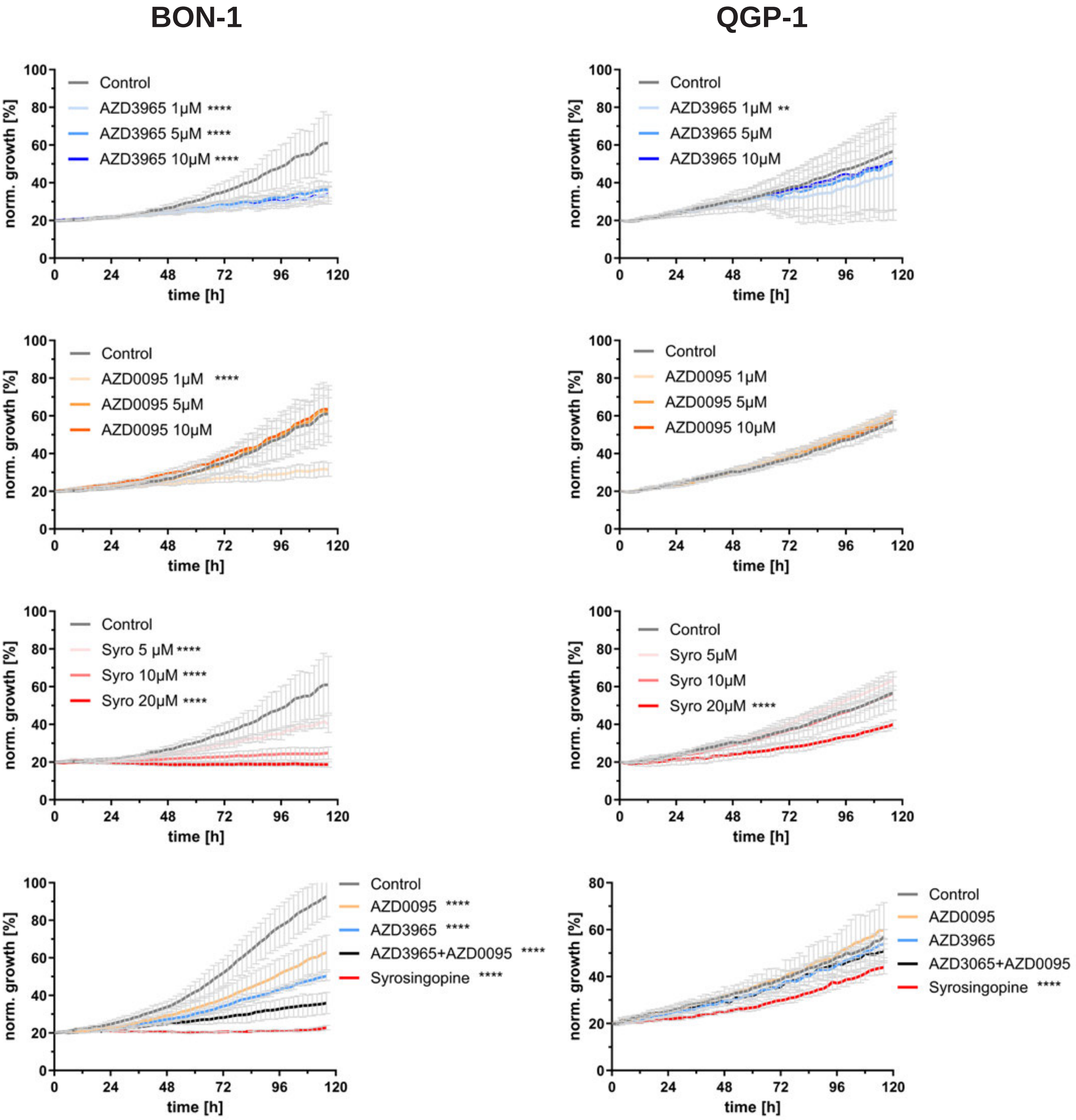

### Suppl Figure 6

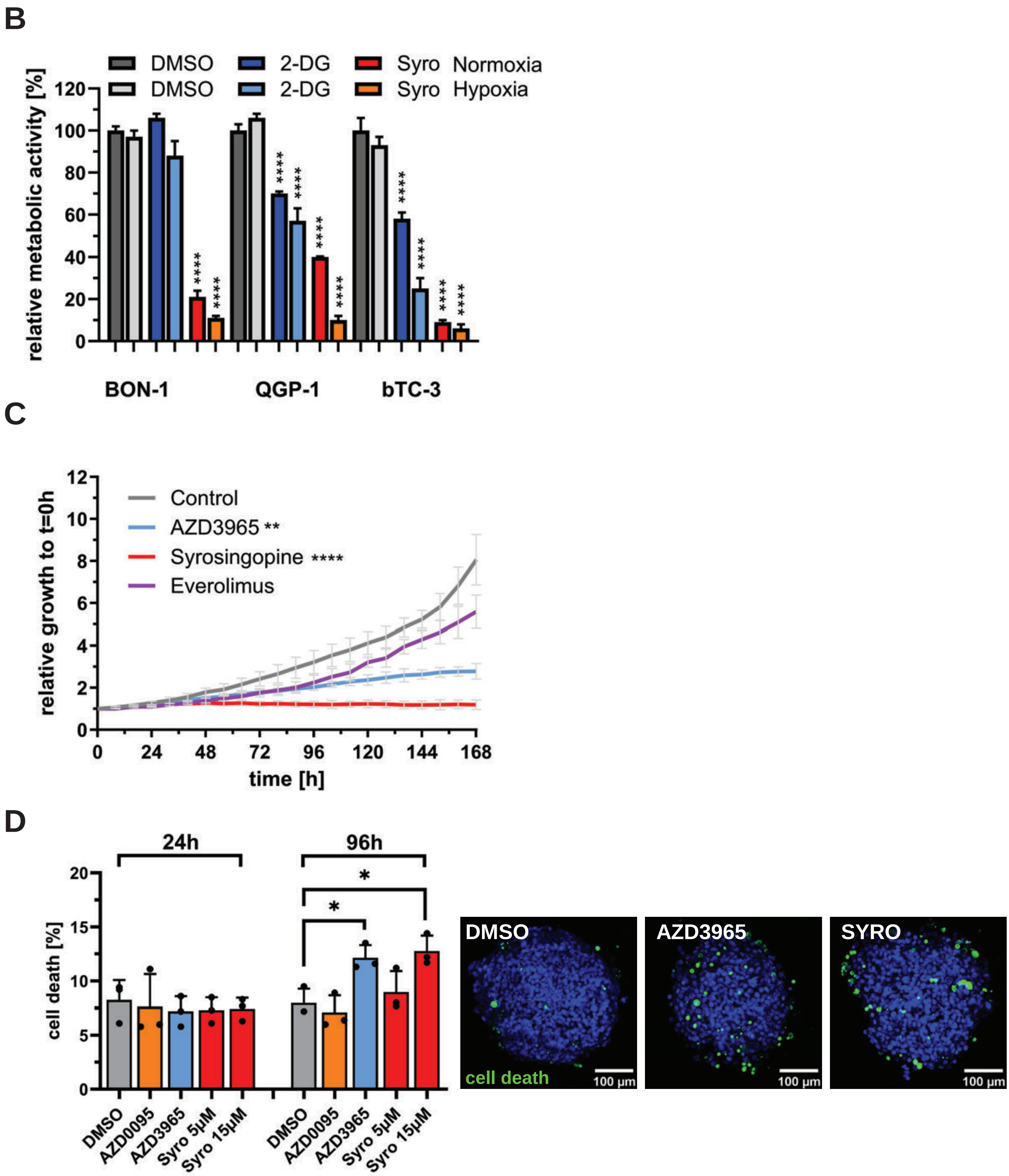

### Suppl Figure 6

E

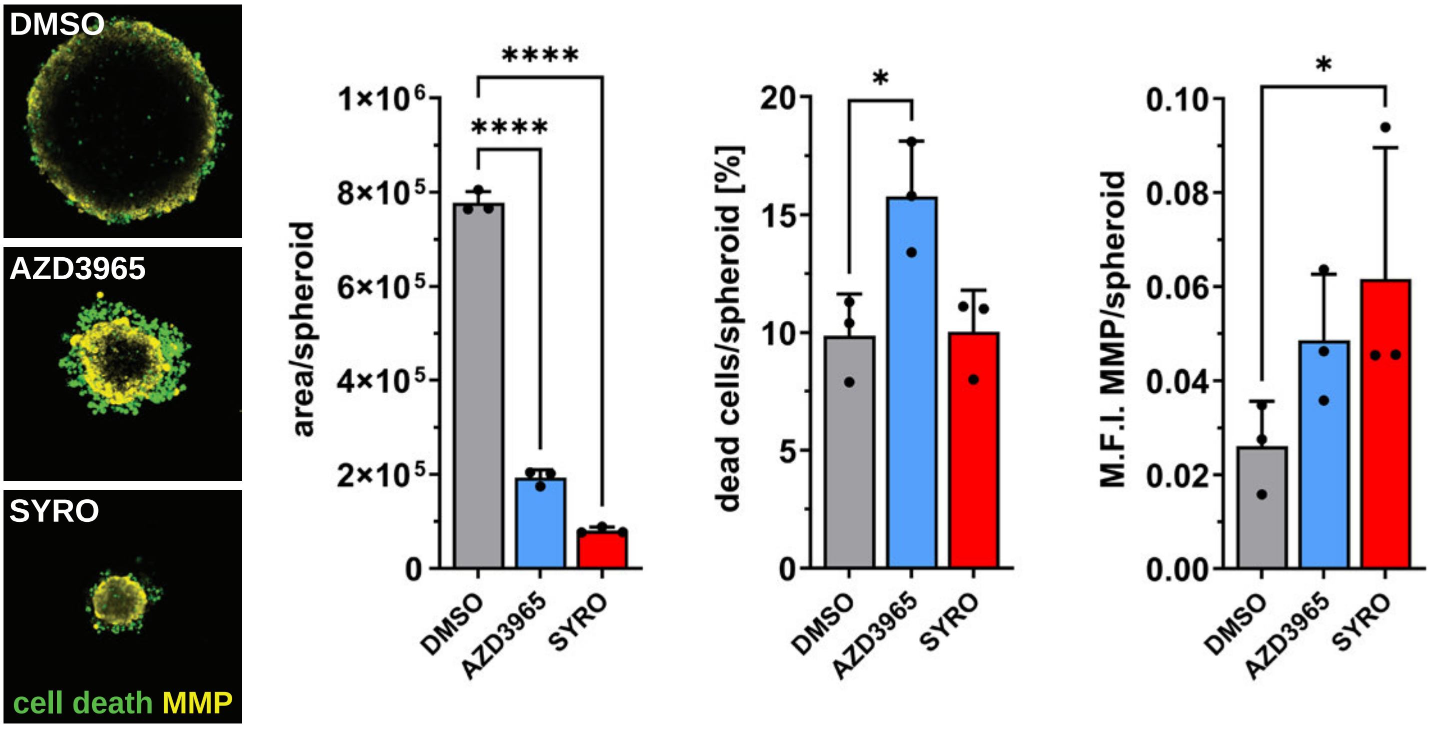

### Suppl Figure 7

## A

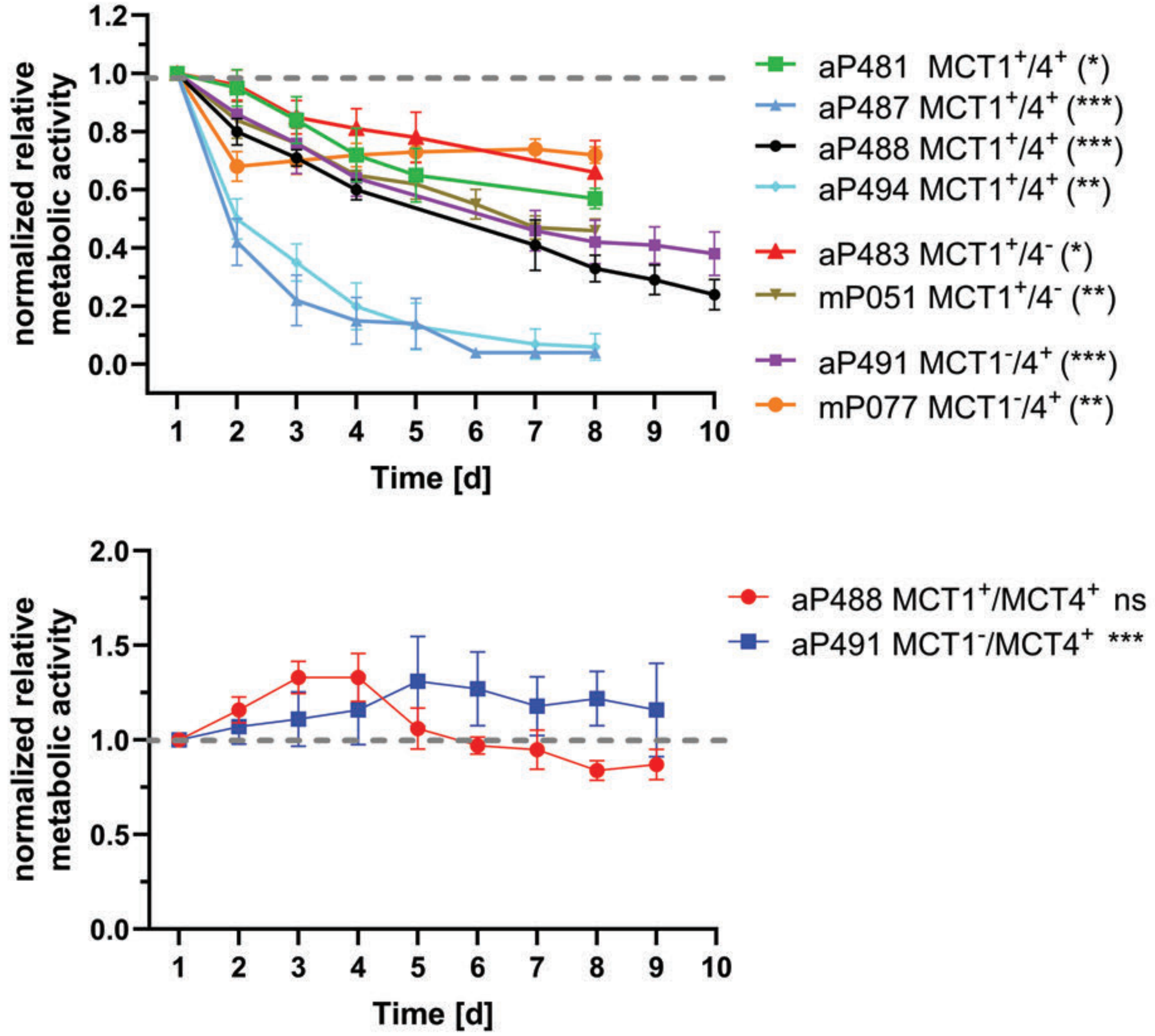


## B

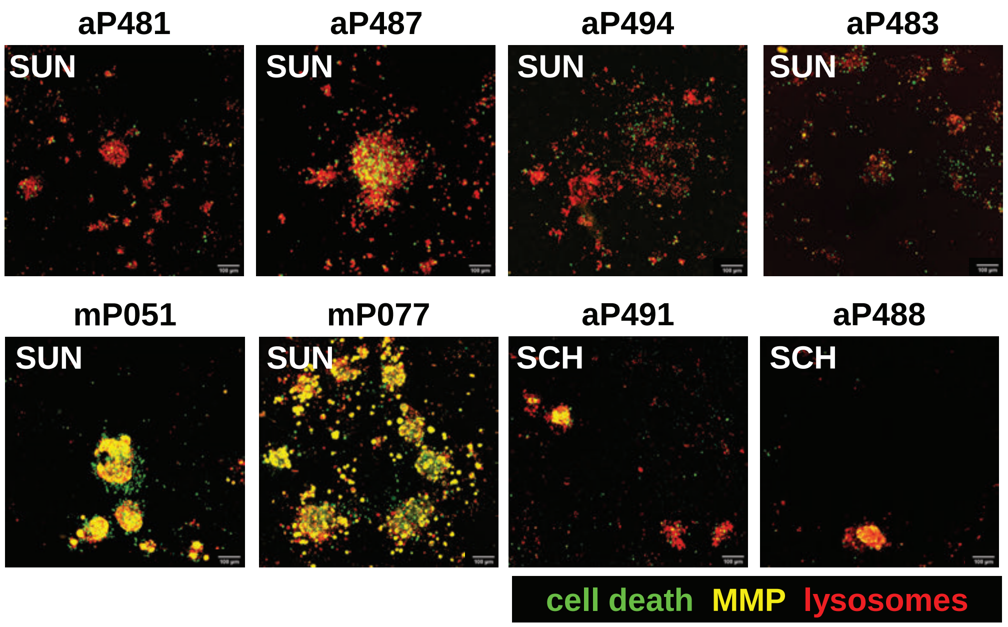
